# Cyclin E1 protein is stabilized in *BRCA1* mutated breast cancers leading to synergy between CDK2 and PARP inhibitors

**DOI:** 10.1101/2020.01.29.911883

**Authors:** Diar Aziz, Neil Portman, Kristine J. Fernandez, Christine Lee, Sarah Alexandrou, Alba Llop-Guevara, Aliza Yong, Ashleigh Wilkinson, C. Marcelo Sergio, Danielle Ferraro, Dariush Etemadmoghadam, David Bowtell, kConFab Investigators, Violeta Serra, Paul Waring, Elgene Lim, C. Elizabeth Caldon

## Abstract

Basal-like breast cancers (BLBC) are aggressive breast cancers that respond poorly to targeted therapies and chemotherapies. In order to define therapeutically targetable subsets of BLBC we examined two markers: cyclin E1 and *BRCA1* loss. In high grade serous ovarian cancer (HGSOC) these markers are mutually exclusive, and define therapeutic subsets. We tested the same hypothesis for BLBC.

Using a BLBC cohort enriched for *BRCA1* loss, we identified convergence between *BRCA1* loss and high cyclin E1 expression, in contrast to HGSOC in which *CCNE1* amplification drives increased cyclin E1 gene expression. Instead, *BRCA1* loss stabilized cyclin E1 during the cell cycle. Using siRNA we showed that *BRCA1* loss leads to stabilization of cyclin E1 by reducing phospho-cyclin E1-T62, and conversely the overexpression of *BRCA1* increased phospho-T62. Mutation of cyclin E1-T62 to alanine increased cyclin E1 stability. We showed that tumors with high cyclin E1/*BRCA1* mutation in the BLBC cohort had decreased phospho-T62, supporting this hypothesis.

Since cyclin E1/CDK2 protects cells from DNA damage and cyclin E1 is elevated in *BRCA1* mutant cancers, we hypothesized that CDK2 inhibition would sensitize these cancers to PARP inhibition. CDK2 inhibition induced DNA damage and synergized with PARP inhibitors to reduce cell viability in *BRCA1* mutated cell lines. Treatment of BLBC patient-derived xenograft models with combination PARP and CDK2 inhibition led to tumor regression and increased survival. We conclude that *BRCA1* status and high cyclin E1 have potential as predictive biomarkers to dictate the therapeutic use of combination CDK inhibitors/PARP inhibitors in BLBC.

## INTRODUCTION

Breast cancer with *BRCA1* mutation most often manifests as basal like breast cancer (BLBC) (1), which presents difficulties for treatment as these cancers present at an earlier age, at a high grade and with greater tumor burden. There are currently no targeted therapies routinely used to treat BLBC (2).

*BRCA1* is a central component of the homologous recombination DNA repair pathway, and its loss results in compromised DNA damage repair (3). Alterations to *BRCA1* are important founder mutations for breast cancer (4), and notably, more than 70% of *BRCA1* mutation carriers develop early-onset BLBC based on gene expression profiling (5). *BRCA1* mutation directly drives the basal phenotype, and mice with *p53* and *BRCA1* deletion develop mammary tumors with basal-like characteristics (6) while intact *BRCA1* represses the transcription of basal cytokeratins (7).

A previous report identified that BLBCs from patients with germline *BRCA1* mutation was associated with high cyclin E1 protein expression (8). Cyclin E1 is a cell cycle regulatory protein whose gain can promote both increased proliferation and genomic instability in cancer cells, and is frequently elevated in BLBC (9). Perplexingly, in high grade serous ovarian cancer (HGSOC) cyclin E1 amplification and *BRCA1/2* mutation are mutually exclusive, presumably because both aberrations drive genomic instability and together they precipitate lethal genomic damage (10–12).

We recently described two subsets of HGSOC, one where cyclin E1 gene amplification and *BRCA1* mutation were mutually exclusive, and another where high cyclin E1 protein expression was due to post–transcriptional deregulation rather than gene amplification, and was often concurrent with *BRCA1/2* mutation (12). Cyclin E1 protein stability is regulated by a multi-step process of specific phosphorylation and ubiquitination, leading to its cyclic expression and turnover (13). Key regulators in the turnover of cyclin E1, such as the ubiquitin ligase component FBXW7 and the deubiquitinase USP28, are frequently dysregulated in cancer (13–15) leading to altered stability of the cyclin E1 protein.

In this study, we examined whether *BRCA1* loss and cyclin E1 gain occurred concurrently or independently in breast cancer. We also explored the mechanisms underpinning high cyclin E1 expression in *BRCA1* mutated breast cancer including gene amplification and protein stability. Finally we tested the hypothesis that disruption of cyclin E1/CDK2 function would sensitize *BRCA1*-mutant cells to PARP inhibition by enhancing synthetic lethality.

## METHODS

### Patient demographics and tumor samples

The Kathleen Cuningham Foundation Consortium for research into Familial Breast cancer (kConFab; http://www.kconfab.org) cohort comprised 308 breast cancer samples (16). Ethics board approval was obtained for patients’ recruitment, sample collection and research studies. Written informed consent was obtained from all participants.

From four tissue microarrays (TMAs), 222 samples had sufficient tumor tissue for immunohistochemistry (IHC). Patient and tumor characteristics are shown in Table 1. Tumors were classified based on estrogen receptor, progesterone receptor, HER2 status and mutation status of *BRCA1*, *BRCA2*, *CHEK1* and *PALB2*. CK5/14 and/or EGFR positive tumors (N=75, 31.6%) were classified as BLBC, while CK5/14 and EGFR negative tumors were classified as non-basal like breast cancer (NBLBC).

**Table 1:**
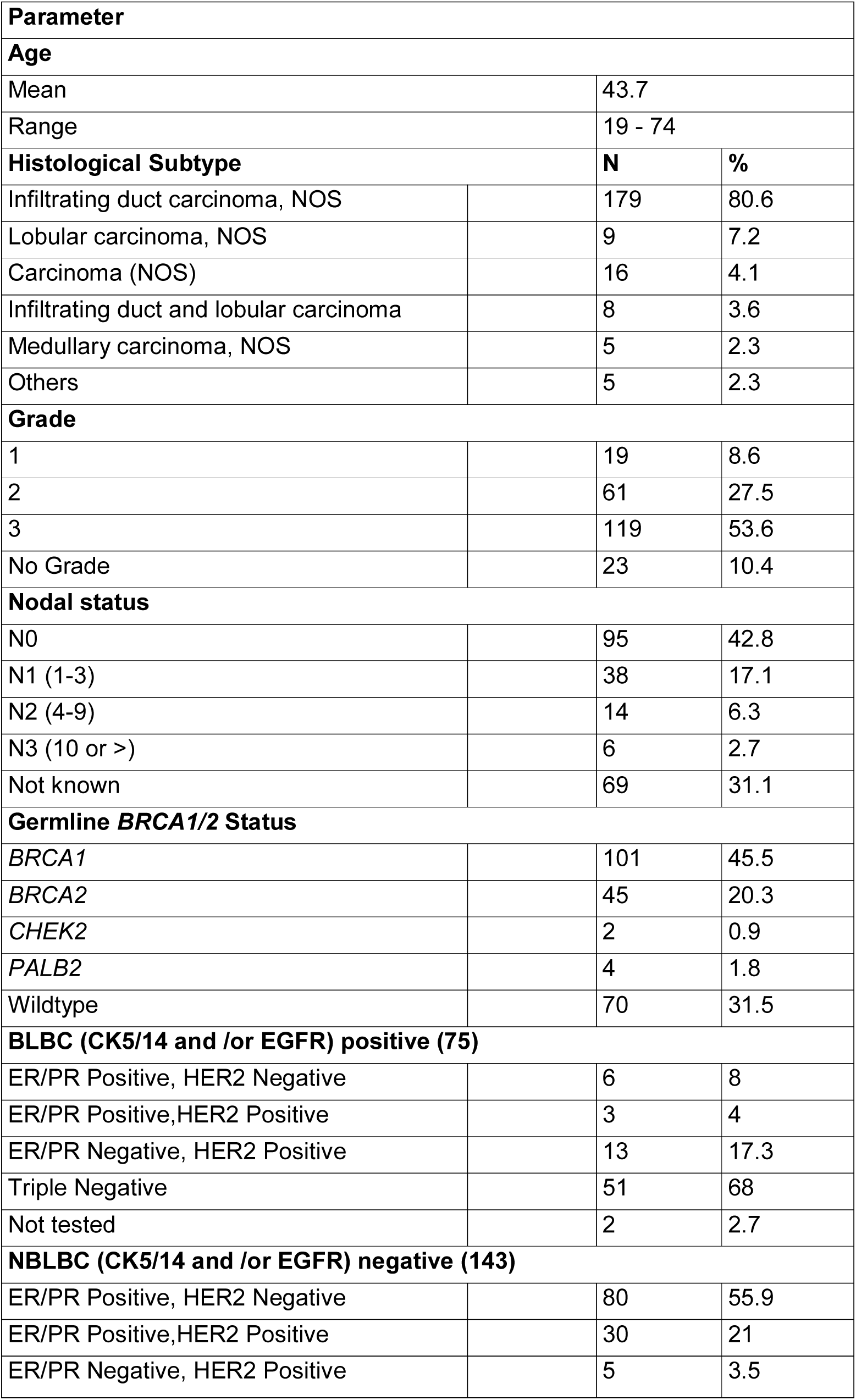
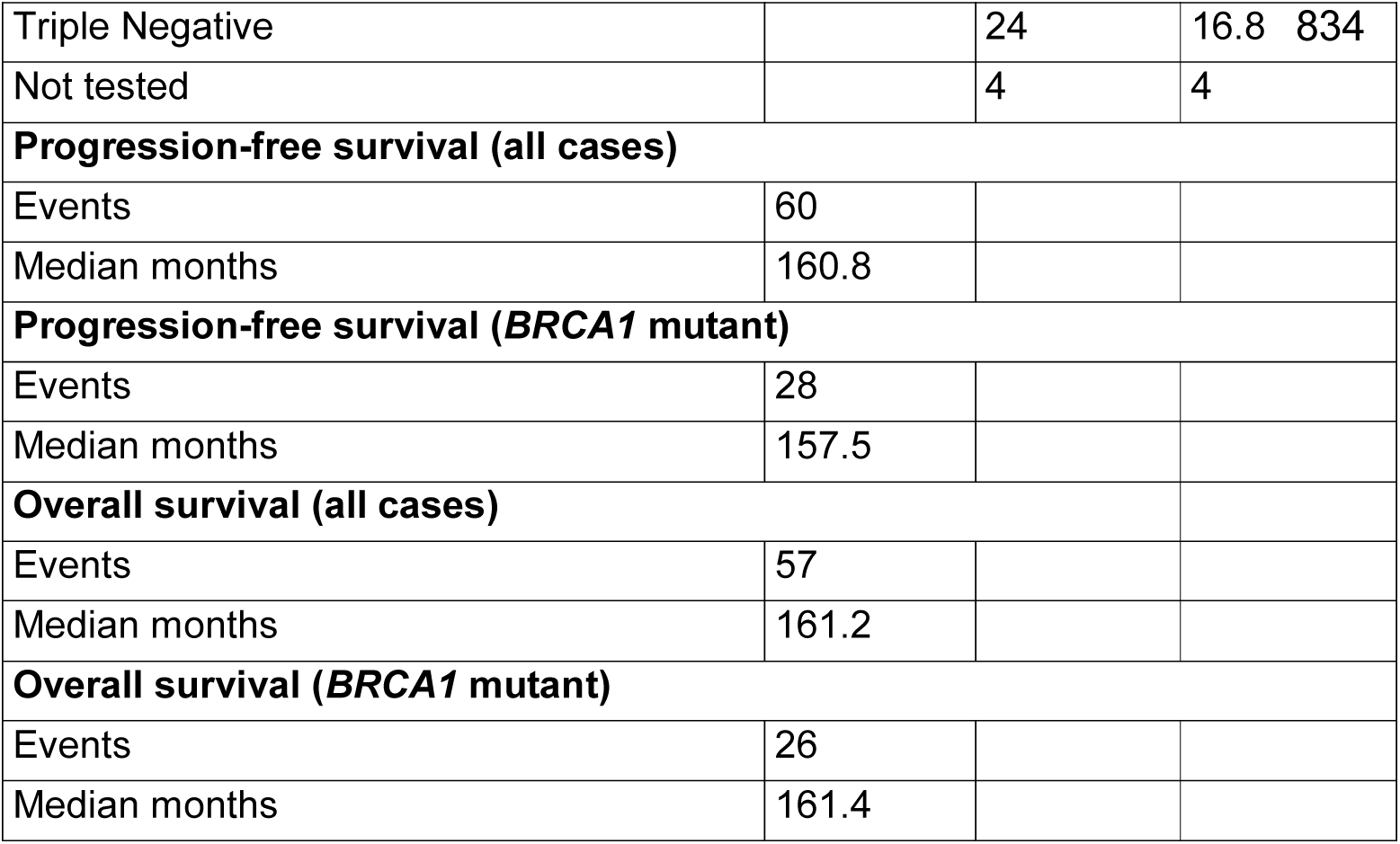
Patient Characteristics.

| **Parameter** |  |  |  |
| --- | --- | --- | --- |
| **Age** |  |  |  |
| Mean |  | 43.7 |  |
| Range |  | 19 - 74 |  |
| **Histological Subtype** |  | **N** | **%** |
| Infiltrating duct carcinoma, NOS |  | 179 | 80.6 |
| Lobular carcinoma, NOS |  | 9 | 7.2 |
| Carcinoma (NOS) |  | 16 | 4.1 |
| Infiltrating duct and lobular carcinoma |  | 8 | 3.6 |
| Medullary carcinoma, NOS |  | 5 | 2.3 |
| Others |  | 5 | 2.3 |
| **Grade** |  |  |  |
| 1 |  | 19 | 8.6 |
| 2 |  | 61 | 27.5 |
| 3 |  | 119 | 53.6 |
| No Grade |  | 23 | 10.4 |
| **Nodal status** |  |  |  |
| N0 |  | 95 | 42.8 |
| N1 (1-3) |  | 38 | 17.1 |
| N2 (4-9) |  | 14 | 6.3 |
| N3 (10 or >) |  | 6 | 2.7 |
| Not known |  | 69 | 31.1 |
| **Germline *BRCA1/2* Status** |  |  |  |
| *BRCA1* |  | 101 | 45.5 |
| *BRCA2* |  | 45 | 20.3 |
| *CHEK2* |  | 2 | 0.9 |
| *PALB2* |  | 4 | 1.8 |
| Wildtype |  | 70 | 31.5 |
| **BLBC (CK5/14 and /or EGFR) positive (75)** |  |  |  |
| ER/PR Positive, HER2 Negative |  | 6 | 8 |
| ER/PR Positive,HER2 Positive |  | 3 | 4 |
| ER/PR Negative, HER2 Positive |  | 13 | 17.3 |
| Triple Negative |  | 51 | 68 |
| Not tested |  | 2 | 2.7 |
| **NBLBC (CK5/14 and /or EGFR) negative (143)** |  |  |  |
| ER/PR Positive, HER2 Negative |  | 80 | 55.9 |
| ER/PR Positive,HER2 Positive |  | 30 | 21 |
| ER/PR Negative, HER2 Positive |  | 5 | 3.5 |

|  |  |  |  |  |
| --- | --- | --- | --- | --- |
| Triple Negative |  | 24 | 16.8 | 834 |
| Not tested |  | 4 | 4 |  |
| **Progression-free survival (all cases)** |  |  |  |  |
| Events | 60 |  |  |  |
| Median months | 160.8 |  |  |  |
| **Progression-free survival (*BRCA1* mutant)** |  |  |  |  |
| Events | 28 |  |  |  |
| Median months | 157.5 |  |  |  |
| **Overall survival (all cases)** |  |  |  |  |
| Events | 57 |  |  |  |
| Median months | 161.2 |  |  |  |
| **Overall survival (*BRCA1* mutant)** |  |  |  |  |
| Events | 26 |  |  |  |
| Median months | 161.4 |  |  |  |

### Dual-colour In Situ Hybridization (ISH) assay for detection of the 19q12 locus amplification

A 19q12 DNP ISH probe that covers the coding sequences of the *CCNE1* and adjacent *URI1* genes and an insulin receptor (INSR) DIG ISH probe, a surrogate reference located on chromosome 19p13.2, were provided by Ventana Medical Systems (Tucson, AZ), and optimized for use on the Ventana ULTRA™ platform as previously described (12), and detailed in the supplementary methods and supplementary figure S1.

### Cyclin E1, FBXW7, USP28 and Phospho-cyclin E1 (T62) IHC

Previously optimized cyclin E1 mouse monoclonal (HE12) (Santa Cruz Biotechnology, CA), FBXW7 rabbit monoclonal (SP-237) (Spring Bioscience, CA) and the USP28 rabbit polyclonal (HPA006778) (Sigma Aldrich) antibody staining were performed using the Ventana Bench Mark ULTRA™ automated staining platform and the Optiview™ Detection kit (12). Optimisation of the cyclin E1 T62 antibody is described in the supplementary methods and supplementary figure S2. All proteins were assessed on nuclear staining using a 0 to 3+ intensity score. Heterogeneous expression was captured using the semi-quantitative H score (17). The distribution of H scores for cyclin E1, phospho-cyclin E1 T62, FBXW7 and USP28 is shown in supplementary figure S3, and determination of the assay cut points for each marker is detailed in supplementary methods.

### Survival Analysis

Kaplan-Meier curves were used to plot overall survival. Assessment of progression free survival was not possible as progression coincided with death in many cases.

### Cell lines and drug treatment

Cell lines were obtained from ATCC and cultured in RPMI1640, 5–10% fetal calf serum (FCS) and insulin (10 μg/ml). All cell lines were authenticated by STR profiling (CellBank Australia) and cultured for less than 6 months after authentication. Cyclin E1 was mutated by site-directed mutagenesis as described (18). MDA-MB-468 cells expressing the ecotropic receptor (19) were infected with pMSCV-IRES-GFP retrovirus expressing cyclin E1 wildtype and mutants as described (20). Subpopulations with graded expression of GFP and cyclin proteins were separated by sterile FACS and matched populations selected based on GFP expression. We expanded cell populations expressing a similar intensity of GFP signal from each cell line, and confirmed expression using western blotting.

Cells were treated with the following drugs resuspended in DSMO: rucaparib (Selleck), paclitaxel (Selleck). CYC065 was provided by Cyclacel Ltd, Dundee, UK.

### Cell proliferation and survival analysis

Survival assays were performed on MDA-MB-468 cells set up at 15,000 per 6 cm dishes in 50% conditioned medium. Paclitaxel (0, 2.6, 2.8 nM) was added every 6-7 days for 3 weeks. Colonies were fixed with trichloroacetic acid (TCA 16%), and stained with 10% Diff Quik Stain 2 (Lab Aids). Quantification was done with ImageJ and the ColonyArea plugin (21).

Metabolic rate was assessed by Alamar Blue (Invitrogen) to determine IC_50_ doses. Synergy assays were performed on indicated cell lines in 96 well plates. The concentration of each drug was increased linearly along each axis of the plate, creating a drug matrix of the different concentrations. The highest concentration of each drug was IC_80_, followed by dilutions of 1/2, 1/4, 1/8, 1/16, and no drug. Cell viability was measured after five days using Alamar Blue. Drug synergy was analysed with Combenefit using the BLISS algorithm (22).

### siRNA transfection

Gene-specific siRNAs to *BRCA1* (On-Target Plus siRNAs J-003461-11-0005 and J-003461-12-0005); *CDK2* (J-003236-11, J-003236-12, J-003236-13, J-003236-14); CDK9 (J-003243-9-0002, J-003243-10-0002, J-003243-11-0002, J-003243-12-0002) and controls [On-Target Plus siCONTROLs (D-001810-10, D-001810-1-4)]; 7 siGENOME Nontargeting siRNA #2 (D-001210-02) were purchased from Dharmacon and transfections carried out as described previously (23).

### Western blot analysis

Cell lysates were extracted and separated on 4-12% Bis-Tris polyacrylamide gels (Invitrogen) as described (24).

Primary antibodies were BRCA1 (#9010, Cell Signalling Technology), USP28 (EPR4249(2), Abcam), CDK2 (M2, Santa Cruz), cyclin E1 (HE12, Santa Cruz), cyclin E2 (EP454Y, Abcam), Mcl-1 (D35A5, Cell Signaling), β-actin (AC-15; Sigma) and GAPDH (4300; Ambion).

### Gene expression analysis

Quantitative reverse transcriptase PCR (qRT-PCR) used inventoried TaqMan probes BRCA1 (Hs01556191_m1), cyclin E1 (Hs00180319_m1) and human RPLPO (4326314E; Applied Biosystems). Analysis was performed as previously described using the ΔΔCT method (25).

### Flow cytometry

S-phase percentages were measured by flow cytometric analysis of propidium iodide stained, ethanol fixed cells. Cell cycle specific expression of endogenous cyclin E1 and V5-tagged cyclin E1 constructs were assessed by flow cytometry as described (26), with further details provided in the Supplementary Data.

### Comet Assay

The alkaline comet assay was performed using the Trevigen Kit (Maryland, USA) according to the manufacturer’s guidelines. HCC1937 cells were seeded in a 6 well plate and treated with CYC065 (Cyclacel) or CVT313 (Thermofisher) at the calculated IC_5_, IC_20_ or IC_50_ dose for 5 days, or treated with 10, 20 or 50nM CDK2 siRNA or CDK9 siRNA, or 50nM non targeting siRNA for 72 hours. Slides were imaged with a fluorescence microscope (Leica DM5500) and analysed with ImageJ OpenComet software (v1.3.1, (27)).

### TCGA datasets

Breast cancer datasets were downloaded via cBioPortal (8) and the BLBC subset identified from PAM50 definitions from TCGA 2015 (28).

### Patient-derived breast cancer xenograft (PDX) models

All *in vivo* experiments, procedures and endpoints were approved by the Garvan Institute of Medical Research Animal Ethics Committee (protocol 18/26) or the VHIO Animal Ethics Committee (protocol 17/42). PDX *BRCA1* R1443* (PDX 11-26) was derived from a metastatic triple negative breast cancer (29), and PDX *BRCA1* 2080delA (PDX124) is basal on PAM50 classification and ER/PR negative (30). At surgery, 3-4 mm^3^ sections of tumor tissue were implanted into the 4^th^ inguinal mammary gland of female NOD-SCID-IL2γR^-/-^ (NSG) mice. PDX *BRCA1* 2080delA animals were implanted subcutaneously and supplemented with 1 μmol/L estradiol (Sigma) in the drinking water. Tumor growth was assessed twice weekly by calliper measurement and mice were randomized to treatment arms when tumors reached 150-250 mm^3^ (using the formula: *width^2^ x length x 0.5*). Vehicle (4% DMSO, 30% PEG-300), 50 mg/kg olaparib and 25mg/kg CYC065 were administered by oral gavage 5 days a week. Mice were treated for 60 days or until tumor volume reached 1000 mm^3^.

### PDX IHC and quantification

Tumor tissue was fixed in 10% neutral buffered formalin and embedded in paraffin, before being sectioned (4uM thick) and stained using the Bond RX Automated Stainer (Leica Biosystems). Heat induced antigen retrieval was performed at pH 9 (Bond Epitope Retrieval solution 2, Leica Biosystems), 100°C for 30mins, before Υ-H2AX antibody incubation (1:500 Cell Signalling, Clone 20E3) for 60mins. Detection was performed with diaminobenzidine (DAB, Bond Polymer Refine Detection, Leica Biosystems) and slides were counterstained with haematoxylin. Slides were imaged using a slide scanner (AperioCS2, Leica Biosystems), and data were analysed using QuPath (31) as described (32).

### Statistical Analysis

Statistical analysis was performed using Prism Software™ version 7 as indicated for each dataset. Data presented as box and whisker plots includes error bars of minimum to maximum, with mean values indicated. Data are presented as column graphs with mean +/- SEM. All experiments were performed in triplicate, except as indicated.

## RESULTS

### *BRCA1* inactivation associates with high cyclin E1 expression in breast cancer

We examined the KConFab cohort, which is enriched for familial cancer mutations, for co-occurrence of germline *BRCA1* mutation and high cyclin E1 expression. First, we examined cyclin E1 expression by IHC (Figure 1A). High cyclin E1 expression was defined as an H score cut-off of ≥45 based on the overall distribution of cyclin E1 expression (Supplementary Figure S3A), and previous reports (9, 33). This cut-off was also associated with patient outcomes (minimal p value, Supplementary Table 1). Overall, germline *BRCA1* mutated cancers had significantly higher cyclin E1 protein than the *BRCA1* wildtype cases, and tumors with other breast cancer associated germline mutations (*BRCA2*, *PALB2*, or *CHK2)* (Figure 1B). Moreover, a significantly larger proportion of germline *BRCA1* mutant cases (82.2%) had detectable cyclin E1 protein (83/101) compared to only 38% of *BRCA1* wildtype tumors (46/121) (P<0.0001, Fisher Exact test).

**Figure 1:**
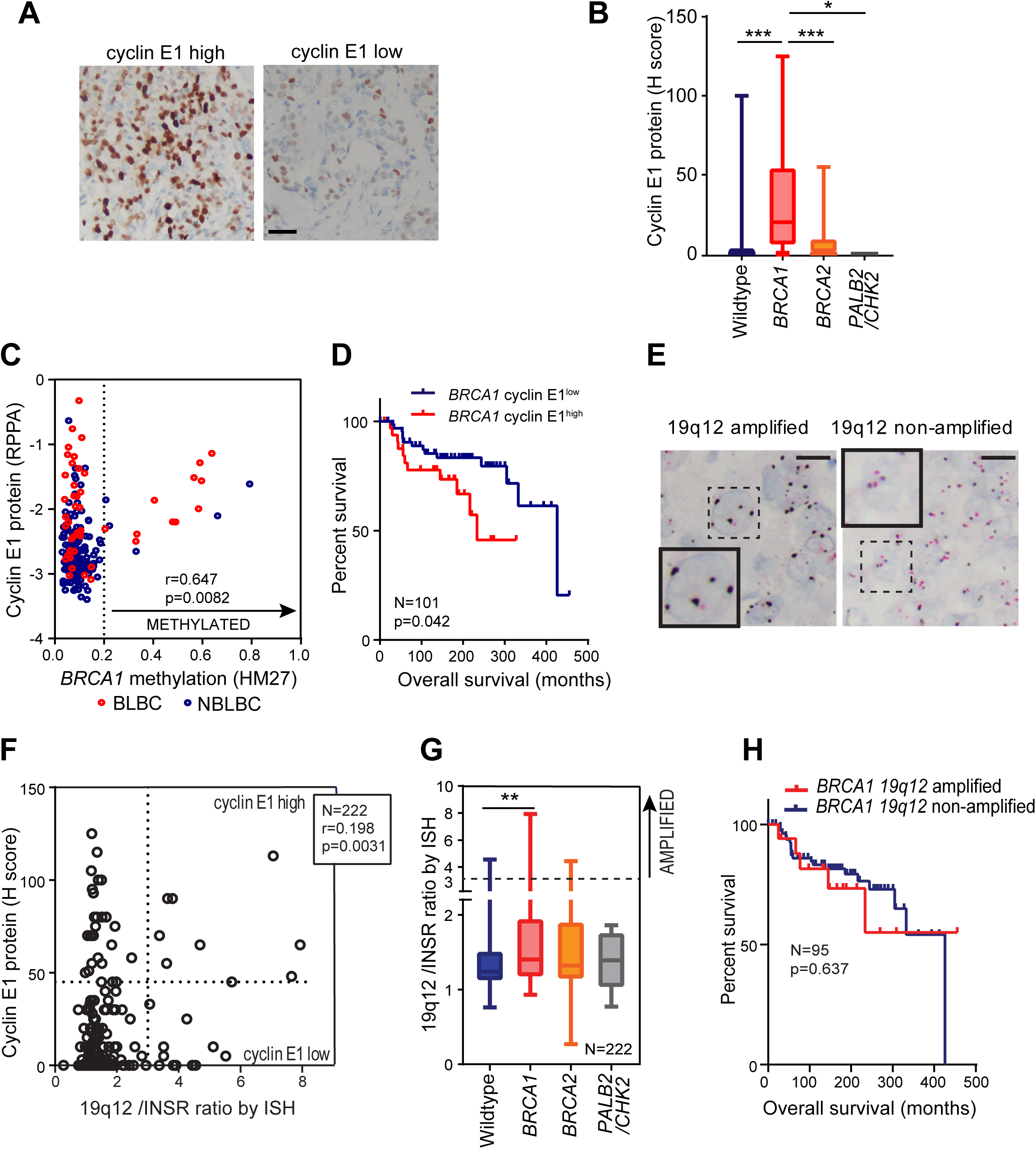
Cyclin E1 is elevated in *BRCA1* deficient cancers, and predicts poor prognosis. **A:** Microscope images of high and low expression of cyclin E1 (IHC). Scale bar is 50µM. **B:** Cyclin E1 protein expression (H score) in wildtype, *BRCA1*, *BRCA2, and PALB2*/*CHEK2* mutated cancers, analysis by one-way ANOVA; * p<0.05; *** p<0.001. **C:** Scatter plot of TCGA breast cancer cohort cyclin E1 protein expression (RPPA) versus *BRCA1* methylation by HM27 array. Dashed line indicates cutoff between methylated and non methylated. Correlation analysis of cyclin E1 protein and *BRCA1* methylation performed across the methylated subset, r=Spearman coefficient. **D:** Kaplan Meier curves of overall survival in the KConfab cohort comparing *BRCA1* mutated cyclin E1 high cases versus *BRCA1* mutated cyclin E1 low cases. **E:** Microscope images of 19q12 non-amplified and 19q12 amplified breast cancer cases (ISH); inset shows representative example of each. Scale bar is 20µM. **F:** Scatter plot of cyclin E1 protein expression versus *CCNE1* (19q12/INSR ratio) amplification status in the KConfab cohort. r=Spearman coefficient. **G:** 19q12/INSR ratio (ISH) cases compared to each of wild type, *BRCA1*, *BRCA2, PALB2*/*CHK2* mutated cancers in the KConfab Cohort, analysed by one way ANOVA. **H:** Kaplan Meier curves of overall survival of *BRCA1* mutated breast cancer comparing 19q12 amplified and non-amplified subsets.

Notably, eight of the germline *BRCA1* wildtype tumors had high cyclin E1. We hypothesized that these may be *BRCA1* methylated since our cohort was selected for familial breast cancers where *BRCA1* methylation is not infrequent (34). Consequently, we examined the relationship between *BRCA1* methylation and cyclin E1 protein expression by interrogating the breast cancer dataset of the TCGA. 241 cases had available data for *BRCA1* methylation and cyclin E1 protein expression. Using a cut-off of 0.2 for methylation (35), we found that *BRCA1* methylation had a significant positive correlation with cyclin E1 protein expression (r=0.647, P=0.0082) (Figure 1C).

Next we examined the association between cyclin E1 expression and overall survival in germline *BRCA1* mutated breast cancers in our cohort. High cyclin E1 expression was associated with a significantly reduced overall survival of patients with *BRCA1* mutation (233.4 vs 426.3 months, P=0.042, HR 0.34, CI 0.130-0.895) (Figure 1D).

### *CCNE1* amplification is not the primary driver of high expression of cyclin E1 in *BRCA1* mutated cancers

The *CCNE1* gene, located on chromosome 19q12, is a frequent site of amplification in cancer. We assessed *CCNE1* amplification by ISH analysis of tissue sections with 19q12 and INSR probes to determine the 19q12/INSR ratio. Representative images of 19q12 non-amplified and 19q12 amplified tumors are shown in Figure 1E. 13.5% (30/222) of tumors in the entire cohort was found to be 19q12 (*CCNE1)* amplified. The correlation between cyclin E1 protein and *CCNE1* gene amplification was poor (r=0.198, P=0.0031, Figure 1F).

Next we assessed whether *CCNE1* amplification and *BRCA1* mutation co-occurred. In contrast to HGSOC (12), 21/101 (20.8%) of *BRCA1* mutant cases had concurrent 19q12 (*CCNE1*) amplification. This was higher than *BRCA1* wildtype cases, where only 6.6% had 19q12 (*CCNE1*) amplification (Figure 1G). Since 19q12 (*CCNE1*) amplification is associated with poor survival in other cancer types we examined its relationship with overall survival in *BRCA1* mutated breast cancer. Unlike high expression of cyclin E1, which is predictive of poor survival, 19q12 (*CCNE1)* amplification had no prognostic value for overall survival in *BRCA1* mutated breast cancer (Figure 1H).

### The cyclin E1 degradation machinery is disrupted in *BRCA1* mutated breast cancers

Since 19q12 status was only poorly predictive of high cyclin E1 expression, we thus investigated other mechanisms that lead to high cyclin E1 expression. One possibility was disruption of the proteasome mediated degradation of cyclin E1, which occurs frequently in cancer (36). Normal cyclin E1 turnover depends upon phosphorylation within two phospho-degrons on the cyclin E1 protein, of which T62 and T380 are crucial phosphorylation sites. The phosphorylated protein is recognized by the FBXW7 module of the SCF^FBXW7^ complex, and ubiquitinated for degradation (13). The deubiquitinase USP28 can remove ubiquitin from cyclin E1 and antagonise FBWX7-mediated degradation (14) (Figure 2A). Disruption of this process, ie loss of cyclin E1 phosphorylation, loss of FBXW7 or gain in USP28, would be expected to increase cyclin E1 stabilization and accumulation.

**Figure 2:**
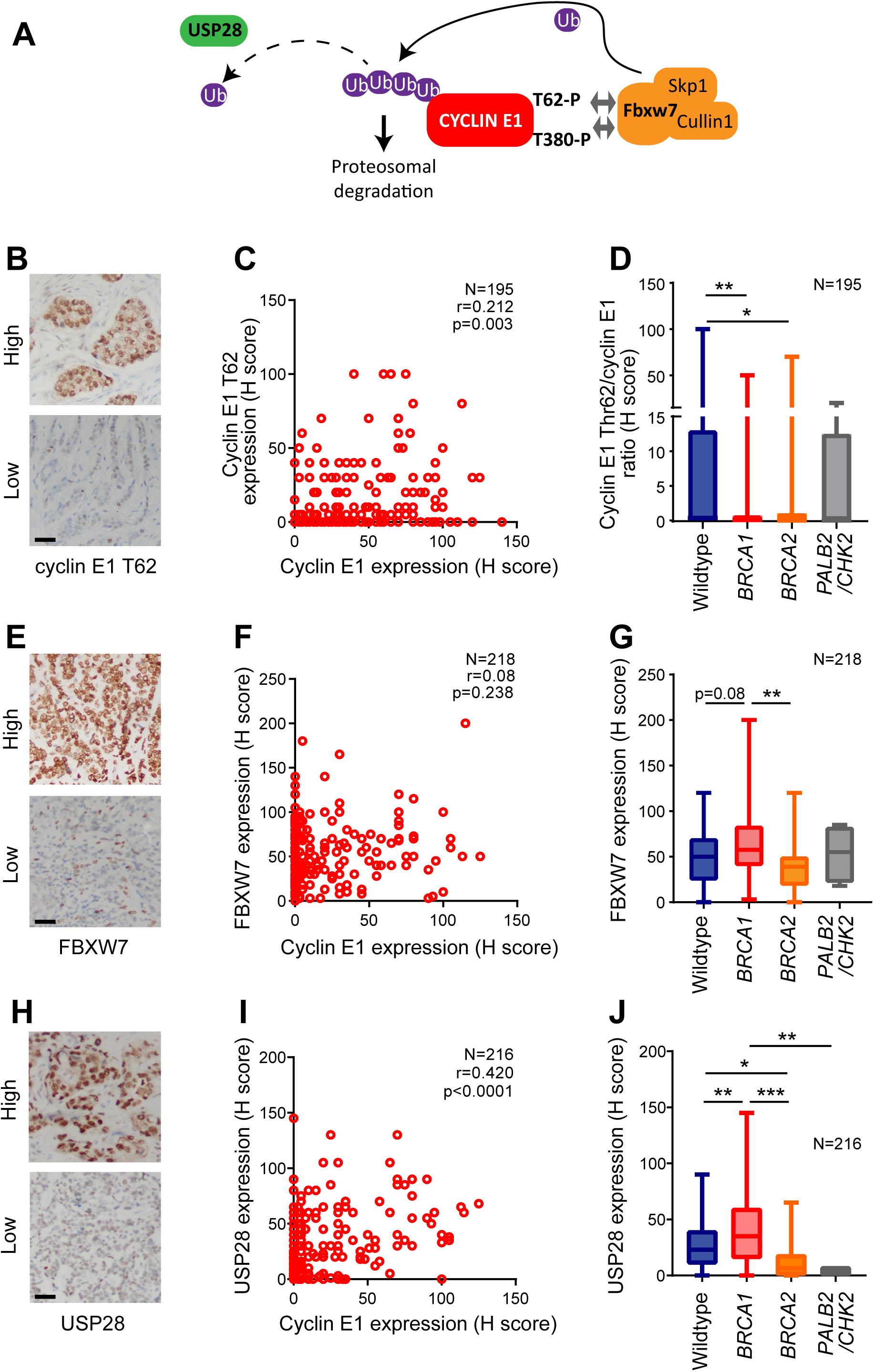
Loss of cyclin E1 T62 phosphorylation and gain of USP28 are associated with *BRCA1* mutation. **A:** Schematic of cyclin E1 turnover. **B:** Microscope images of IHC staining of breast cancer sections with high and low phospho-cyclin E1 T62. Scale bar is 50µM. **C:** Scatter plot of cyclin E1 expression versus phospho-cyclin E1 T62 expression, r=Spearman coefficient. **D:** Phospho-cyclin E1 T62/cyclin E1 ratio of expression in wildtype, *BRCA1* mutated, *BRCA2* mutated*, PALB2*/*CHK2* mutated subsets of the KConfab Cohort, analysed by one way ANOVA; * p<0.05; ** p<0.01. **E:** Microscope images of IHC staining of BC sections with high and low FBXW7. Scale bar is 50µM. **F:** Scatter plot of cyclin E1 expression versus FBXW7 expression, r=Spearman coefficient. **G:** FBXW7 expression in wildtype, *BRCA1* mutated, *BRCA2* mutated*, PALB2*/*CHK2* mutated subsets of the KConfab Cohort, analysed by one way ANOVA; ** p<0.01. **H:** Microscopic images of IHC staining of BC sections with high and low USP28 expression. Scale bar is 50µM. **I:** Scatter plot of cyclin E1 expression versus USP28 expression, r=Spearman coefficient. **J:** USP28 expression in wildtype, *BRCA1* mutated, *BRCA2* mutated*, PALB2*/*CHK2* mutated subsets of the KConfab Cohort, analysed by one way ANOVA; * p<0.05, ** p<0.01, *** p<0.001.

We assessed the cyclin E1 degradation machinery by IHC in our familial breast cancer cohort. We first assessed cyclin E1 T62 phosphorylation in 195 cases (representative images in Figure 2B). We observed only a moderate positive correlation between cyclin E1 T62 phosphorylation and cyclin E1 expression (r=0.212, p=0.003, Spearman), indicating that a proportion of cancers had very low cyclin E1 T62 phosphorylation (Figure 2C). Consequently, we assessed the ratio of cyclin E1 phosphorylation to its absolute expression to determine if phosphorylation was specifically dysregulated in certain subsets of patients. We found that the *BRCA1* and *BRCA2* mutant subsets exhibited a significantly lower T62/cyclin E1 ratio (Figure 2D), indicative of a loss of cyclin E1 phosphorylation in the absence of functional *BRCA1* or *BRCA2*.

There was no correlation between cyclin E1 and FBXW7 found in our cohort (Figure 2E, F). However, *BRCA1* mutated cancers had higher expression compared to the *BRCA2* mutant subset (Figure 2G). In contrast, USP28 expression was moderately correlated with cyclin E1 expression (r=0.420, p<0.0001, Spearman) (Figure 2H, I). USP28 protein expression was significantly higher in the *BRCA1* mutated subset (Figure 2J).

In summary, *BRCA1* mutated breast cancers were characterised by reduced cyclin E1-T62 phosphorylation and elevated USP28 expression. Overall, these data implicate increased cyclin E1 protein stability, rather than gene amplification, as the cause for high cyclin E1 levels observed in *BRCA1* mutated breast cancer.

### *BRCA1* loss leads to cell cycle stabilization of cyclin E1

Since the cyclin E1 degradation machinery was deregulated in *BRCA1* mutant cancers across our cohort, we investigated whether cyclin E1 turnover is dysregulated in cell lines with mutant *BRCA1* or *BRCA1* loss. The BLBC cell line HCC1937 has a homozygous *BRCA1* 5382C* mutation and the triple negative breast cancer (TNBC) cell line MDA-MB-436 has a *BRCA1* homozygous deletion. We compared these to 4 cell lines with wildtype *BRCA1*: BT-20 and MDA-MB-468 (BLBC cell lines), MDA-MB-231 (TNBC), and SkBr3 (HER2 amplified). Cells were analysed for the expression of cyclin E1 during the cell cycle using flow cytometry. BT-20, MDA-MB-468, SkBr3 and MDA-MB-231 cells showed a typical downregulation of cyclin E1 during S phase (Figure 3A), which we quantitated by comparing the expression of cyclin E1 during the second half of S phase versus the first half of S phase (Figure 3B). The *BRCA1* defective cell lines showed significantly diminished down-regulation of cyclin E1 during S phase: in HCC1937 cells the cyclin E1 levels 14 did not decrease in S phase, but instead marginally increased. There was a small decrease in the absolute expression of cyclin E1 during S phase in MDA-MB-436 cells (Figure 3A/B).

**Figure 3:**
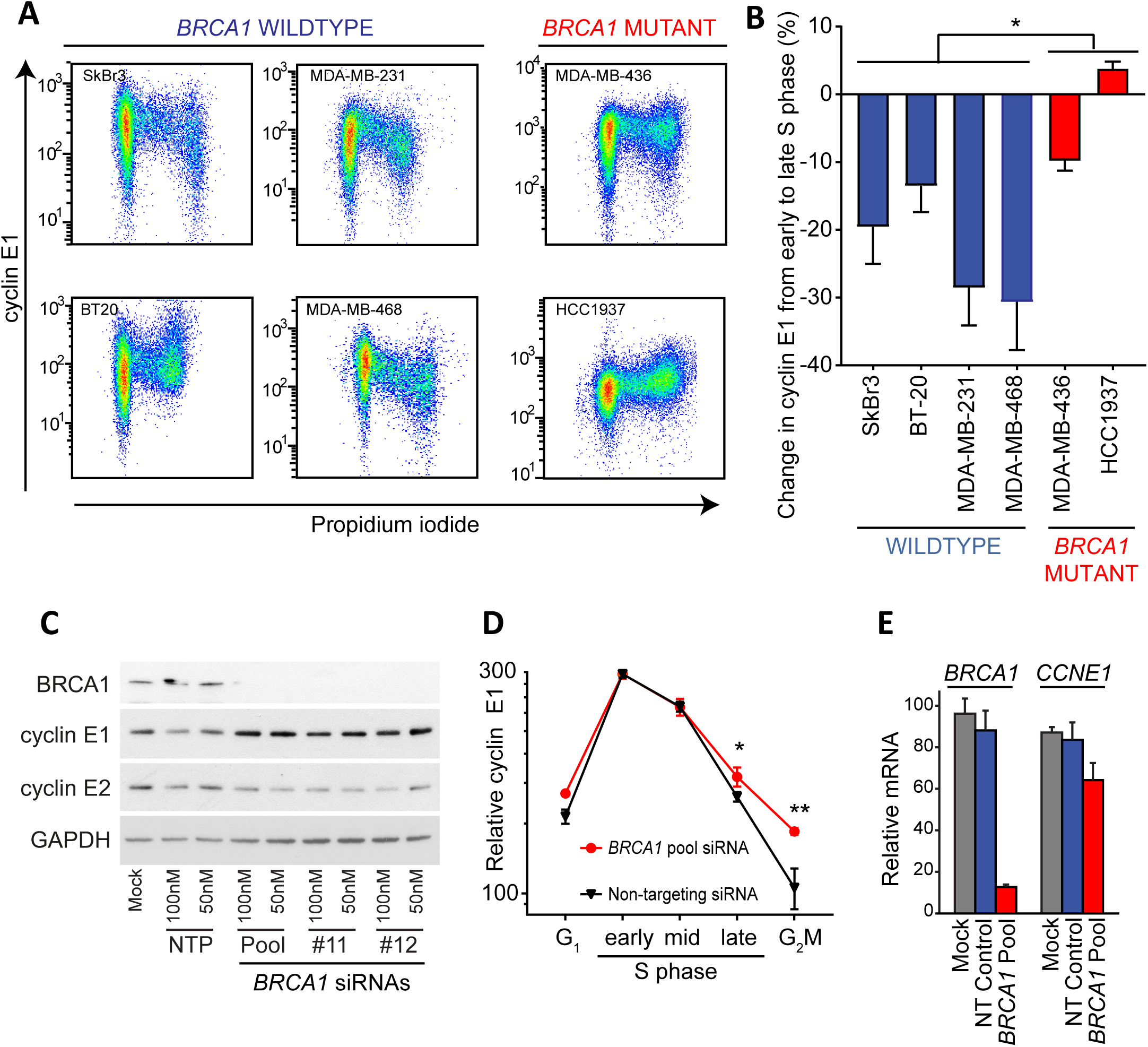
Cyclin E1 protein is stabilized in the absence of functional *BRCA1*. **A:** Breast cancer cell lines (SkBr3, MDA-MB231, BT20 and MDA-MB-468, MDA-MB-436 and HCC1937) were analysed by flow cytometry for intracellular cyclin E1 and DNA content (propidium iodide). **B:** Quantitation of fold change in expression of cyclin E1 from early S phase to late S phase as measured by flow cytometry in (A.). Data is mean +/- SEM, analysed by t-test; * p<0.05. **C:** T-47D cells were treated with siRNAs for 48h (siRNAs: NTP (non-targeting pool) *BRCA1* Pool, #11 and #12), and lysates western blotted for BRCA1, cyclin E1, cyclin E2 and GAPDH. **D:** T-47D cells treated for 48h with BRCA1 pool siRNA, FBXW7 pool siRNA and non-targeting pool siRNA were analysed by flow cytometry for cyclin E1 expression and DNA content (propidium iodide).The geometric mean expression of cyclin E1 at G_1_, early S, mid S, late S and G_2_M phase was quantitated for each treatment, and data is mean +/- SEM; * p<0.05, ** p<0.01. **E:** T-47D cells treated for 48h with BRCA1 pool siRNA and non-targeting pool siRNA were analysed by qRT-PCR for *CCNE1* expression, and data is mean +/- range of duplicate experiments assayed in triplicate.

We next investigated whether knockdown of *BRCA1* was able to recapitulate the cell cycle stabilization of cyclin E1. First, we treated *BRCA1* wildtype T-47D cells, with two different *BRCA1* siRNAs, siRNA#11 and siRNA#12, and a pool of the two siRNAs. Both *BRCA1* siRNAs led to a reproducible increase of cyclin E1 protein (Figure 3C). Notably, this was specific to cyclin E1, and did not affect cyclin E2 protein (Figure 3C). We then specifically assessed whether *BRCA1* knockdown led to changes in the cell cycle expression of cyclin E1. T-47D breast cancer cells were transfected with pooled *BRCA1* siRNA followed by flow cytometry analysis of cells immunoprobed for cyclin E1 and co-stained with propidium iodide. *BRCA1* siRNA treatment led to a significant increase in and prolongation of cyclin E1 protein expression during late S phase and G_2_/M of the cell cycle (Figure 3D). In contrast, *CCNE1* mRNA was not increased following *BRCA1* siRNA exposure (Figure 3E), confirming that the increase in cyclin E1 expression occurs post-transcriptionally.

### *BRCA1* dysregulation specifically alters T62 phosphorylation of cyclin E1, but not USP28 expression

Since cyclin E1 cell cycle expression is dysregulated with *BRCA1* disruption and we had observed both loss of T62 phosphorylation and gain of USP28 in our cohort, we sought to confirm that *BRCA1* loss alters either T62 phosphorylation or USP28 expression to stabilise cyclin E1.

We first tested whether *BRCA1* knockdown stabilized cyclin E1 via upregulation of USP28. siRNA-mediated knockdown of USP28 in MDA-MB-231 cells led to decreased expression of cyclin E1 (Supplementary Figure S4A). In contrast *BRCA1* siRNA led only to downregulation of BRCA1 but did not change USP28 levels (Supplementary Figure S4B). Thus while USP28 is elevated in *BRCA1* mutant cancers, we could not detect its regulation directly downstream of *BRCA1*.

Next we examined whether there was a direct relationship between changes to *BRCA1* expression and cyclin E1 phosphorylation. We exposed T-47D cells to *BRCA1* siRNA to increase cyclin E1 expression (Figure 4A), and subsequently immunoprecipitated the lysates with the phospho-cyclin E1 T62 and phospho-cyclin E1 T380 antibodies to examine the relative abundance of phosphorylation of cyclin E1 at these sites. Cyclin E1 T62 was depleted following *BRCA1* siRNA, in contrast cyclin E1 T380 expression was sustained (Figure 4A). Thus the increased expression of cyclin E1 following depletion of BRCA1 protein was linked directly to cyclin E1 T62 dephosphorylation.

**Figure 4:**
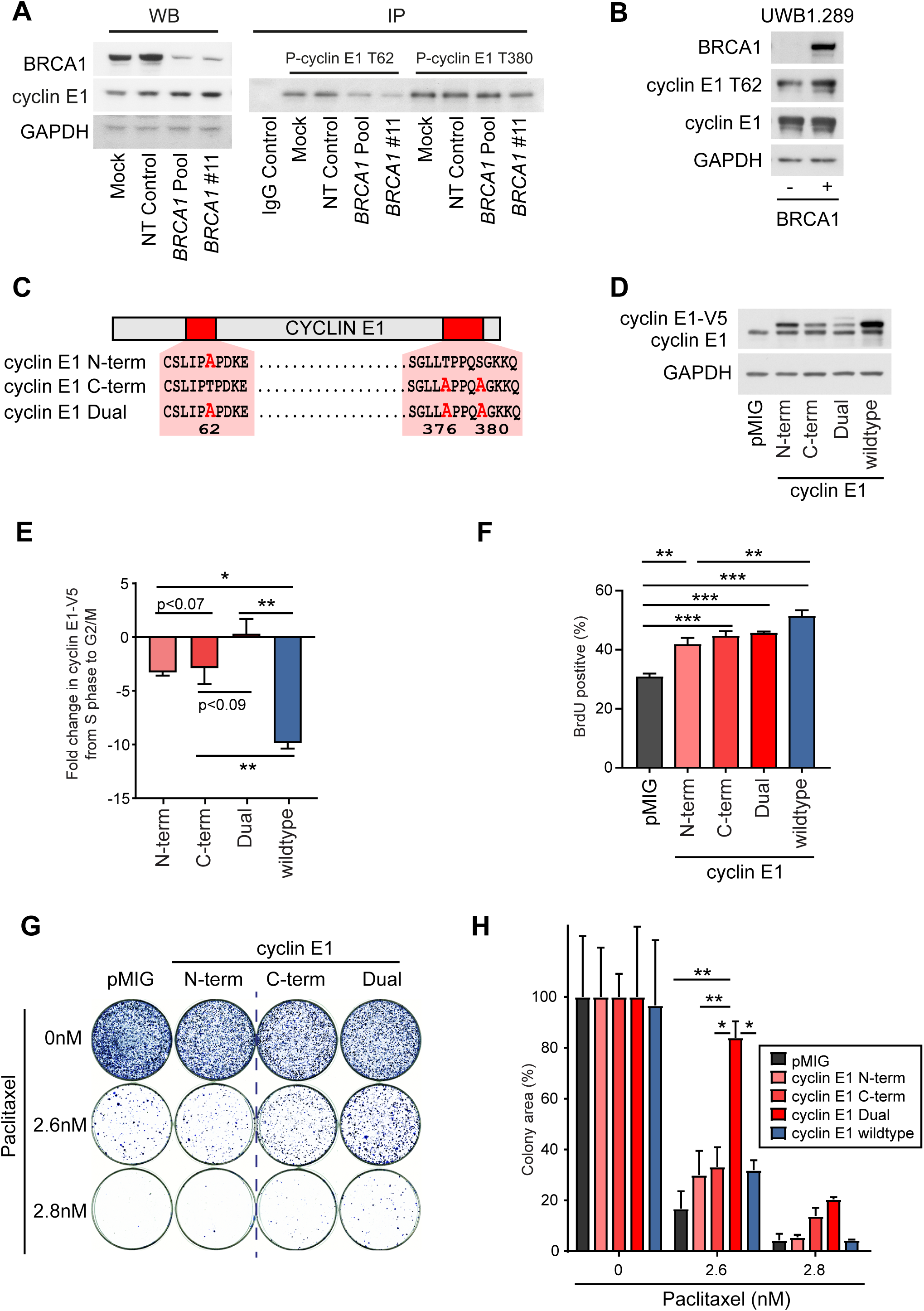
*BRCA1* loss leads to decreased cyclin E1 T62 phosphorylation, which alters protein stability, and contributes to proliferation and cell survival. **A:** T-47D cells were treated with siRNAs for 48h (siRNAs: NTP (non-targeting pool) *BRCA1* Pool, #11), and lysates western blotted for BRCA1, cyclin E1, and GAPDH. Lysates were also immunoprecipitated with phospho-T62-cyclin E1, phospho-T380 cyclin E1 and a rabbit IgG control, and immunoprecipitates western blotted for cyclin E1. **B:** UWB1.289 and UWB1.289 BRCA1 cells were western blotted for BRCA1, phospho-T62 cyclin E1, cyclin E1 and GAPDH. **C:** Schematic of site-directed mutagenesis of phospho-sites of cyclin E1. **D:** MDA-MB-468 cells were retrovirally infected with V5-tagged cyclin E1 constructs (N-term, C-term, Dual, wildtype), sorted by flow cytometry for populations with matched GFP expression, and lysates western blotted for cyclin E1 and GAPDH. **E:** MDA-MB-468 cells expressing cyclin E1 constructs (N-term, C-term, Dual, wildtype) were analysed by flow cytometry for V5-cyclin E1 expression and DNA content (propidium iodide).The geometric mean expression of cyclin E1 at early S and late S phase was quantitated for each treatment, and the fold change from early to late S phase is shown as the mean +/- SEM of triplicate experiments. Data analysed by one-way ANOVA; * p<0.05, ** p<0.01. **F:** MDA-MB-468 cells expressing cyclin E1 constructs (N-term, C-term, Dual, wildtype) were analysed by flow cytometry for BrdU incorporation. Data is the mean +/- SEM of triplicate experiments, analysed by one-way ANOVA; ** p<0.01, *** p<0.001. **G:** MDA-MB-468 cells expressing cyclin E1 constructs (N-term, C-term, Dual, wildtype) were treated with paclitaxel (0nM, 2.6nM, 2.8nM) for 3 weeks, and colony formation detected with Diff Quick Stain 2. **H:** Colony formation was quantitated using the ColonyArea ImageJ plugin from triplicate assays. Data is the mean +/- SEM analysed by two-way ANOVA; * p<0.05, ** p<0.01.

We performed the converse analysis by comparing UWB1.289 ovarian cancer cells (germline *BRCA1* mutation within exon 11 along with deletion of the wild-type allele) to UWB1.289/*BRCA1* cells, which stably re-express *BRCA1* (37), and found that the *BRCA1* restored cell line had higher cyclin E1 T62 expression (Figure 4B).

Since T62 dephosphorylation can increase cyclin E1 protein stability, we analysed the effect of disrupting the phospho-degrons on cyclin E1 by mutating phospho-sites to alanine to mimic the non-phosphorylated state. We performed site-directed mutagenesis within the two phospho-degrons of cyclin E1 (Figure 4C). We created an N-terminal mutant (T62A, designated N-term), a C-terminal mutant (T376A/S380A, designated C-term) and a combined mutant (T62A/T376A/S380A, designated Dual) and stably expressed these in MDA-MB-468 BLBC cells (Figure 4D).

We examined the effect of each mutant on the stability of the cyclin E1 protein by performing flow cytometry for the V5 tag protein during the cell cycle. We measured the fold change in each of the V5 tagged proteins between early and late S phase (Supplementary Figure S5). All three mutants were significantly more stable than the wildtype “high” cyclin E1 protein (Figure 4E). The T62A site in the N-terminus stabilizes the cyclin E1 protein, and particularly in combination with mutation of the C-terminal phospho-sites of cyclin E1.

Next, we examined the effect of each mutant on cell proliferation. Overexpression of cyclin E1 wildtype and each of the cyclin E1 mutants led to a significant increase in BrdU incorporation compared to the vector control (Figure 4F).

Following this we examined whether these mutants were able to alter the survival of cells when treated with paclitaxel, a taxane chemotherapy used to treat BLBC clinically (38). We treated vector control, cyclin E1 wildtype, and cyclin E1 mutant cells with paclitaxel, and monitored survival by colony forming assay after 3-4 weeks. Only the Dual mutant cells demonstrated significantly increased colony counts compared to wildtype cyclin E1 overexpression (Figure 4G/4H).

Overall, *BRCA1* loss led to decreased cyclin E1 T62 phosphorylation, which in turn can increase cyclin E1 protein stability and percentage of cells in S phase. Cyclin E1 T62 was also critical in combination with other cyclin E1 phosphorylation sites to increase cell survival in the presence of paclitaxel.

### Synergistic targeting of cells with high cyclin E1 and *BRCA1* mutation

Our data showing that *BRCA1* loss has a direct role in sustaining elevated cyclin E1 protein levels during S phase, supporting the rationale of co-targeting these proteins. It has been demonstrated that *BRCA1* deficiency leads to susceptibility to inhibition of poly (ADP-ribose) polymerase (PARP), whereas cyclin E1 activates the therapeutically targetable kinase CDK2. However, CDK2 also has important roles in DNA repair (39), leading to increased sensitivity of *BRCA1/2* mutant cancers to CDK2 inhibitors (40). We thus hypothesized that treating *BRCA1* mutant cancers with a combination of CDK2 and PARP inhibitors would be synergistic due to the simultaneous blockade of cyclin E1 dependent proliferation and exacerbated synthetic lethality from PARP inhibitors due to the additional DNA damage resulting from CDK2 inhibition.

First, we tested whether CDK2 inhibition induces DNA damage, by treating *BRCA1* mutant HCC1937 cells with two CDK2 inhibitors, CYC065, and CVT313. After establishing a dose response curve for CYC065 and CVT313 (Supplementary Figure S6A and S6B), we identified induction of DNA damage using the alkaline Comet assay, which detects both double strand and single strand DNA breaks (Figure 5A/B). CYC065 targets CDK2, CDK5 and CDK9, but with the highest specificity to CDK2. CDK5 has negligible expression in HCC1937 cells (41). We subsequently confirmed that DNA damage was occurring via CDK2 action by performing comet assays after treatment with CDK2 and CDK9 siRNA treatment. CDK2 siRNA treatment led to an increase in tail moment detection after 72 hours of exposure at all doses (Figure 5C). In contrast, there was no effect on DNA damage following CDK9 siRNA exposure (Figure 5D).

**Figure 5:**
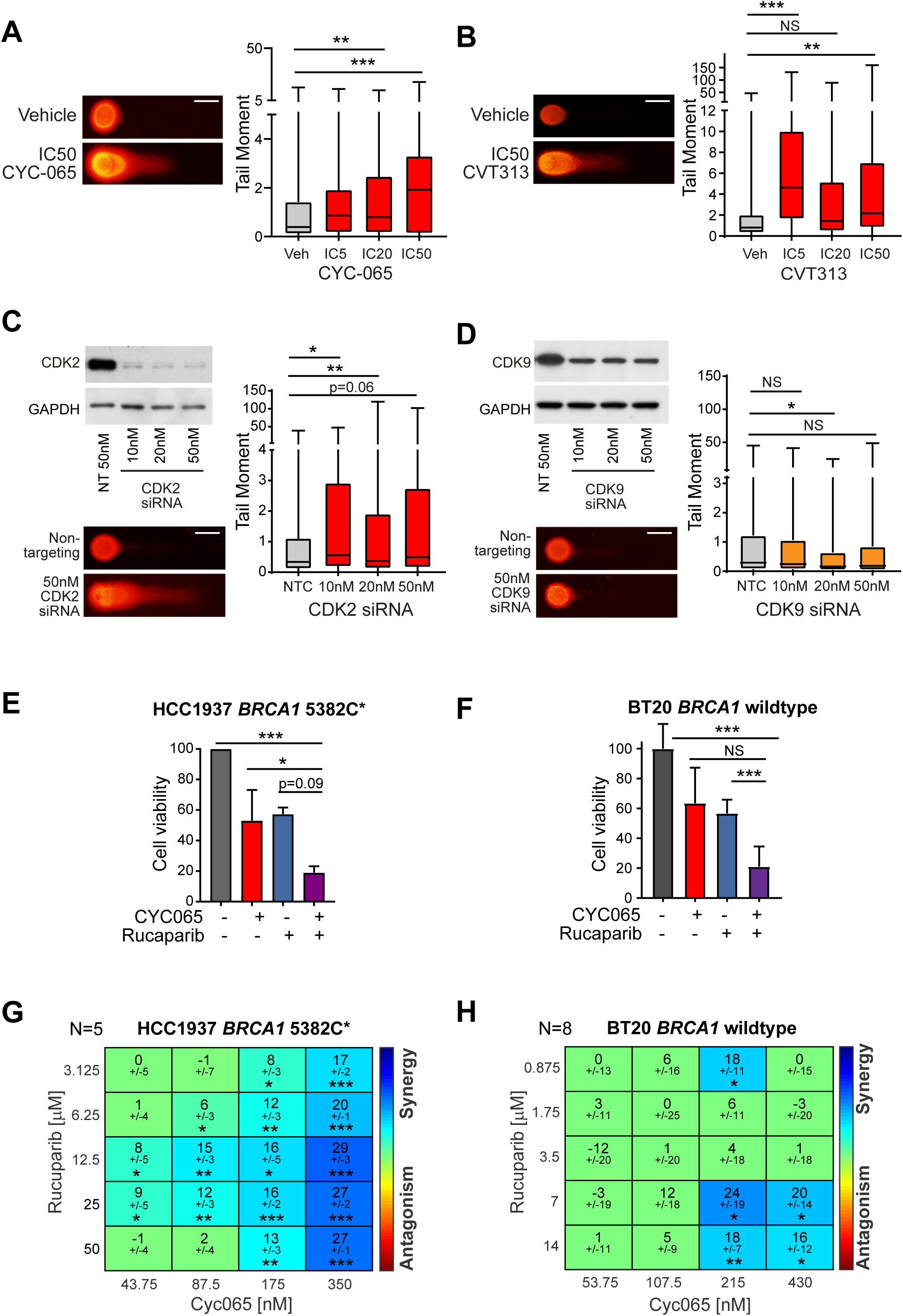
CDK2 inhibition induces DNA damage to synergise with PARP inhibition in *BRCA1* defective breast cancer cells. **A:** HCC1937 cells treated with CYC065 or vehicle for 5 days were analysed by alkaline Comet assay, 90-300 tails quantitated/treatment in triplicate experiments. Data analysed by one-way ANOVA; ** p<0.01, *** p<0.001. Representative images shown, scale bar is 50μM. **B:** HCC1937 cells treated with CVT313 or vehicle for 5 days were analysed by alkaline Comet assay, 90-300 tails quantitated/treatment in triplicate experiments. Data analysed by one-way ANOVA; ** p<0.01, *** p<0.001. Representative images shown, scale bar is 50μM. **C:** HCC1937 cells treated with *CDK2* siRNA or Non-targeting Pool siRNA for 72h were analysed by alkaline Comet assay, 190-250 tails quantitated/treatment. Quadruplicate experiments analysed by one-way ANOVA; * p<0.05, ** p<0.01. Representative images shown, scale bar is 50μM. **D:** HCC1937 cells treated with *CDK9* siRNA or Non-targeting Pool siRNA for 72h were analysed by alkaline Comet assay, 190-250 tails quantitated/treatment. Triplicate experiments analysed by one-way ANOVA; * p<0.05. Representative images shown, scale bar is 50μM. **E:** HCC1937 cells were treated with IC50 doses of CYC065, rucaparib, or CYC065 + rucaparib for 5 days, as well as vehicle control, and cell viability measured by Alamar Blue. Triplicate experiments were analysed by one way ANOVA; * p<0.05, *** p<0.001. **F:** BT20 cells were treated with IC50 doses 33 of CYC065, rucaparib, or CYC065 + rucaparib for 5 days, as well as vehicle control, and cell viability measured by Alamar Blue. Triplicate experiments were analysed by one way ANOVA; *** p<0.001. **G:** HCC1937 cells were treated with doses of CYC065 and rucaparib for 5 days, and viability measured by Alamar Blue. Synergy analysis was performed by BLISS, where blue indicates synergy, and red indicates antagonism. Data is pooled from 5 replicates; * p<0.05, ** p<0.01, *** p<0.001. **H:** BT20 cells were treated with doses of CYC065 and rucaparib for 5 days, and viability measured by Alamar Blue. Synergy analysis was performed by BLISS, where blue indicates synergy, and red indicates antagonism. Data is pooled from 8 replicates; * p<0.05, ** p<0.01.

Next, we examined the effect of combining CDK2 inhibition with PARP inhibition. We treated a *BRCA1* wildtype (BT20) and *BRCA1* mutant cell line (HCC1937) with CYC065 and rucaparib (a PARP inhibitor). After first establishing dose response curves (Supplementary data S6A/S6C), we exposed the cells to IC_50_ doses of CYC065 and rucaparib. The combination was significantly more effective than either drug used as a single agent in the *BRCA1* mutant and wildtype cell lines (Figure 5E-F). There was a significant synergy demonstrated with BLISS analysis between the two drugs at intersecting dose curves of CYC065 and rucaparib (Figure 5G-H). This occurred over a greater dose range in the *BRCA1* mutant compared to the wildtype cells, where the effect was predominantly additive.

### Combination olaparib and CYC065 treatment leads to tumor regression *in vivo*

We tested *in vivo* efficacy of the combination therapy in two PDXs of BLBC origin with pathogenic *BRCA1* alterations: *BRCA1* R1443* mutation and truncating *BRCA1* 2080delA mutation. Following tumor implantation and expansion, each model was treated with daily gavage of 50mg/kg olaparib (a PARP inhibitor) and 25mg/kg CYC065 (sub-optimal doses selected for combination testing) (Figure 6A). In the *BRCA1* R1443* mutant PDX, single agent olaparib led to reduced tumor burden and increased overall survival (Figure 6B-D). There was no single agent response to CYC065. By contrast, all olaparib and CYC065 combination treated tumors regressed to below the starting tumor volume, and the host mice survived until the experimental endpoint (Figure 6B-D).

**Figure 6:**
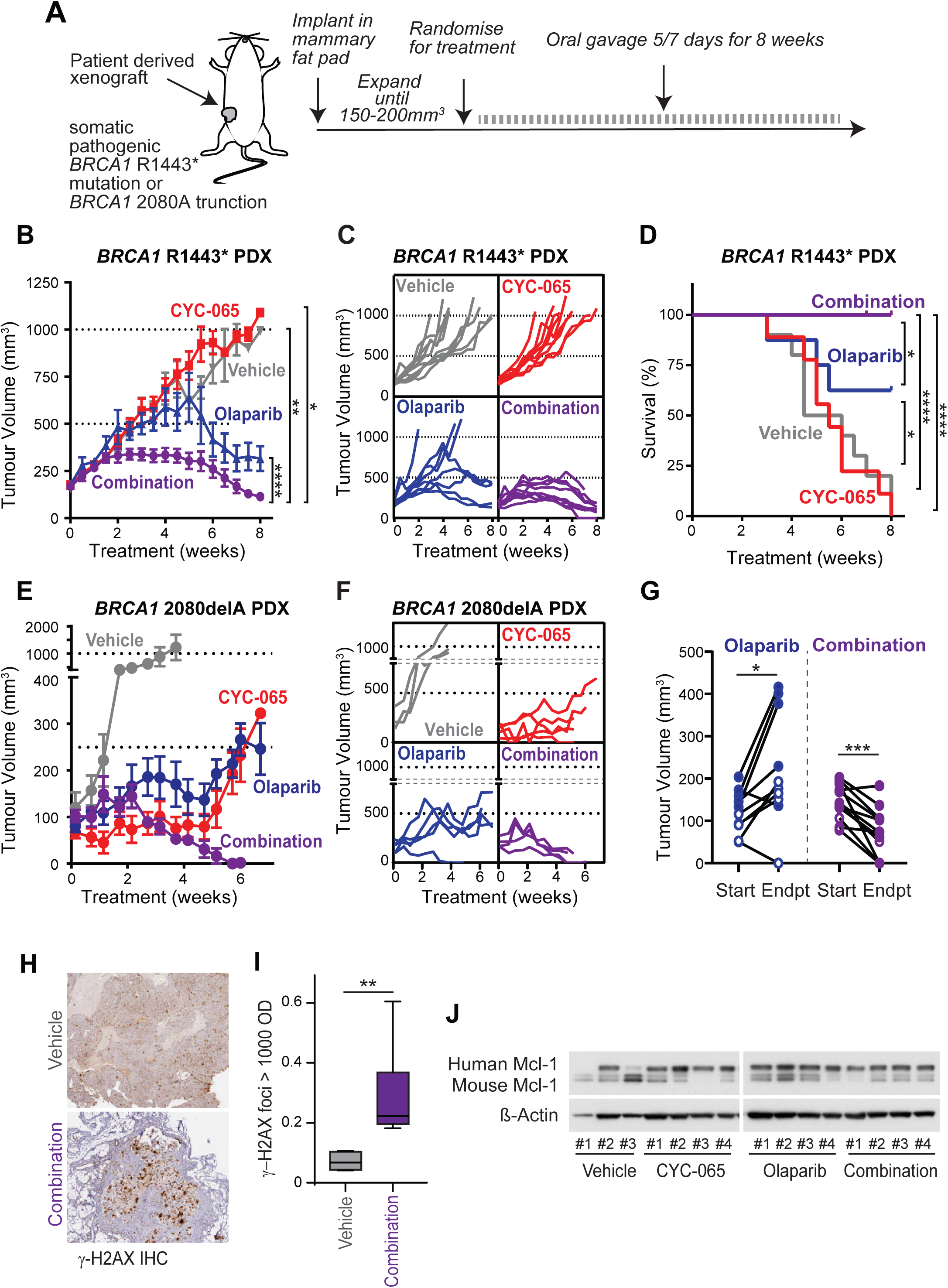
The combination of CDK2 inhibition with PARP inhibition leads to tumor regression in *BRCA1* mutant PDX models. **A:** Schematic for drug administration to BLBC PDX models with *BRCA1* mutation. **B:** BLBC *BRCA1* R1443* PDX model was treated with vehicle (grey, n=10), CYC065 25mg/kg (red, n=9), olaparib 25mg/kg (blue, n=8) or the combination of CYC065 and olaparib (purple, n=9). Tumor volume was measured with calipers for 8 weeks. Data analysed by repeated measures one-way ANOVA; * p<0.05, ** p<0.01, **** p<0.0001. **C.** Growth kinetics of individual *BRCA1* R1443* PDX tumors with therapy. **D.** Kaplan-Meier survival curves of (B.), statistical differences between curves estiumated by the Logrank (Mantel Cox) test; * p<0.05, **** p<0.0001. **E.** BLBC *BRCA1* 2080delA PDX model was treated with vehicle (grey, n=3), CYC065 25mg/kg (red, n=5), olaparib 25mg/kg (blue, n=5) or the combination of CYC065 and olaparib (purple, n=4). Tumor volume was measured with calipers for 7 weeks. **F.** Growth kinetics of individual *BRCA1* 2080delA PDX tumors with therapy. **G.** The change in tumor volume in both *BRCA1* R1443* and *BRCA1* 2080delA PDX models between start of treatment and ethical endpoint for Vehicle treated cohort. Endpoint for *BRCA1* R1443* PDX was 60 days. Endpoint for *BRCA1* 2080delA PDX was 33 days. Data analysed by two-tailed paired t-test; * p<0.05, *** p<0.001. **H.** γ-H2AX IHC of Vehicle and olaparib/CYC065 treated tumors of the *BRCA1* R1443* PDX models. **I.** Quantitation of high intensity γ-H2AX foci in Vehicle and olaparib/CYC065 treated tumors, analysed by two-sided t-test. **J.** Western blots for Mcl-1 and GAPDH in lysates from Vehicle (n=3), CYC065 (n=4), olaparib (n=4) and combination CYC065/olaparib (n=4) treatment.

The *BRCA1* 2080delA model has high expression of cyclin E1, probably driven in part by *CCNE1* copy number gain (1.31x from exome sequencing). In this model we found that both single agent olaparib and CYC065 were effective, leading to a significant reduction in tumor volume. However, by the 60 day endpoint of the experiment, several of the tumors treated with either olaparib or CYC065 had begun to grow in the presence of therapy. In contrast, the combination therapy was highly effective and resulted in tumor regression at the experimental endpoint (Figure 6E, F).

Next, we compared the effect of combination therapy to olaparib alone across the two models. Olaparib therapy resulted in smaller tumors compared to controls, but they were significantly larger than the starting volume (1.85x larger, P <0.013). In contrast, treatment with the combination therapy led to the reduction in size of all but one tumor across the two cohorts (0.51x smaller, p<0.0005; Figure 6G). Finally, since CYC065 acts via CDK2 and CDK9 inhibition, we examined the specific induction of DNA damage to determine if there was evidence for inhibition of CDK2 in the combination treated tumors. Tumors at endpoint for vehicle showed diffuse H2AX staining across the entire tumor, whereas combination treated tumors showed intense H2AX foci and entire cells positive for H2AX (Figure 6H), with significantly more intense staining in the combination treated tumors (Figure 6I). We also examined a canonical marker of inhibition of CDK9, the downregulation of cell survival protein, Mcl-1. Mcl-1 expression was maintained or higher in CYC065, olaparib, and olaparib/CYC065 treated tumors than in vehicle treated tumors (Figure 6J).

## DISCUSSION

The PARP inhibitors olaparib and talazoparib were recently FDA approved for use as monotherapy in patients with metastatic germline *BRCA1/2*-mutated breast cancer based on significant improvement in progression free survival compared to chemotherapy (42). However, the utility of PARP inhibitors in BLBC is limited as clinical trials did not show an improvement in overall survival, and partial and complete responses were infrequent. Combinations with chemotherapy can be limited by myelosuppression (43). Consequently, there is a compelling unmet clinical need to identify targeted therapies that enhance the lethality of PARP inhibitors without precipitating intolerable side-effects.

We here find that *BRCA1* loss reduces the turnover of cyclin E1 thereby increasing proliferation and survival, providing a new therapeutic opportunity to enhance the synthetic lethality of PARP inhibitors by co-targeting the cyclin E1/CDK2 axis. We have used CYC065, a CDK2/CDK5/CDK9 inhibitor which has successfully completed a First-in-Human Phase I clinical trial, and is continuing clinical development in both solid tumors and haematological malignancies. Our studies identify that CDK2 inhibitors work specifically through CDK2 to induce DNA damage *in vitro*. We demonstrate high efficacy of combination CYC065 and olaparib *in vivo* to induce tumor regression. Olaparib as a single agent was effective in our PDX models, but several individual tumors were shown to escape therapy, and overall tumor burden was increased by the experimental endpoints. We show no tumors escaped inhibition with combination treatment, and almost all tumors regressed. We note that the *in vivo* models may also have benefited from the additional inhibition of CDK9 via CYC065, although with the conditions used we do not observe the downregulation of Mcl-1 which is normally associated with CDK9 inhibition (44) (Figure 6J). Pan-CDK inhibitors that target CDK1 or CDK12 have been demonstrated in pre-clinical TNBC models to result in HR deficiency and induce synthetic lethality in combination with PARP inhibitors (29). This has led to the pan-CDK inhibitor dinaciclib being trialed clinically in combination with the PARP inhibitor veliparib in a patient cohort with TNBC (Clinical Trials Gov reference NCT01434316). Our data indicates that these patients may similarly derived benefit from synthetic lethality between CDK2 inhibition and PARP inhibition.

We found that cyclin E1 is stabilized in *BRCA1* mutated breast tumors via reduced phosphorylation on cyclin E1 Threonine 62, and high cyclin E1 is associated with decreased overall survival for patients with *BRCA1* mutation. Cyclin E1 T62 phosphorylation was originally believed to be of lesser importance in the turnover of cyclin E1, but our work here, and that of others (45), shows that it can have potentially strong effects in tumorigenesis. We find that T62A mutation is sufficient to increase cyclin E1 stability and BrdU incorporation, and that T62A mutation contributes to cell survival in combination with mutation of the other major phospho-sites of the protein. Mutation of cyclin E1 T74A and cyclin E1 T393A (equivalent to human cyclin E1 T62 and T395) in a mouse model led to much higher cyclin E1 levels in hematopoietic and epithelial cells compared to T393A mutation alone, as well as hematopoietic neoplasia (45). Delayed mammary gland involution after pregnancy was also observed exclusively in the presence of the T74A mutation (45), highlighting its likely importance for breast tumorigenesis.

The kinase responsible for T62 phosphorylation has not been identified, though it is hypothesized to be a CDK2 auto-phosphorylation site based on a loose consensus sequence for CDK2 around the T62 site, and the timing of T62 phosphorylation early in G_1_ phase soon after partnering with CDK2 (46). Consequently, increased T62 auto-phosphorylation may be the result of a direct physical interaction between *BRCA1* and cyclin E1/CDK2 (47) or through downstream effectors of *BRCA1* action. We observed that *BRCA1* mutation-mediated stabilization only occurred for the cyclin E1 protein, but not the closely related ortholog cyclin E2 (Figure 3C). This is despite the phospho-T62 site being conserved between the two proteins (48). However, unique to cyclin E1 is an upstream GSK3-β consensus site at S58 which is hypothesized to require T62 phosphorylation in order to be phosphorylated (49).

In summary, we have found that CDK2 inhibition may sensitize *BRCA1* mutant breast cancer cells to PARP inhibitors. *BRCA1* mutation most commonly associates with the aggressive BLBC subtype, and thus the presence of *BRCA1* mutation in concert with the BLBC phenotype would suggest combination CDK2 and PARP inhibition as an effective therapeutic strategy. As low levels of *BRCA1* and *BRCA1* methylation are also very common to BLBC (50), and our data demonstrates elevated cyclin E1 in the *BRCA1* methylated BLBC, a rational ongoing area of investigation is CDK2 inhibition to sensitize *BRCA1* methylated or deficient cancers to PARP inhibitors.

## ACKNOWLEDGMENTS

We wish to thank Heather Thorne, Eveline Niedermayr, Sharon Guo, all the kConFab research nurses and staff, the heads and staff of the Family Cancer Clinics, and the Clinical Follow Up Study (which has received funding from the NHMRC, the National Breast Cancer Foundation, Cancer Australia, and the National Institute of Health (USA)) for their contributions to this resource, and the many families who contribute to kConFab. kConFab is supported by a grant from the National Breast Cancer Foundation, and previously by the National Health and Medical Research Council (NHMRC), the Queensland Cancer Fund, the Cancer Councils of New South Wales, Victoria, Tasmania and South Australia, and the Cancer Foundation of Western Australia.

D.A. is a recipient of the Higher Committee of Education of Iraq Scholarship. V.S. is supported by Spanish Instituto de Salud Carlos III (ISCIII) funding, an initiative of the Spanish Ministry of Economy and Innovation partially supported by European Regional Development FEDER Funds: Miguel Servet [CP14/00228], FIS [PI17/01080] and Transcan-2 TH4RESPONS (AC15/00063). A.L-G. is supported by a PERIS fellowship from the Departament de Salut, Generalitat de Catalunya [SLT002/16/00477]. EL is a National Breast Cancer Foundation Endowed Chair and supported by Love Your Sister. C.E.C. is supported by a National Breast Cancer Foundation Career Development Fellowship (EC17-002), and is grateful for the support of the Dr Lee McCormick Edwards Charitable Foundation.

---

**Supplementary Figure S1:**
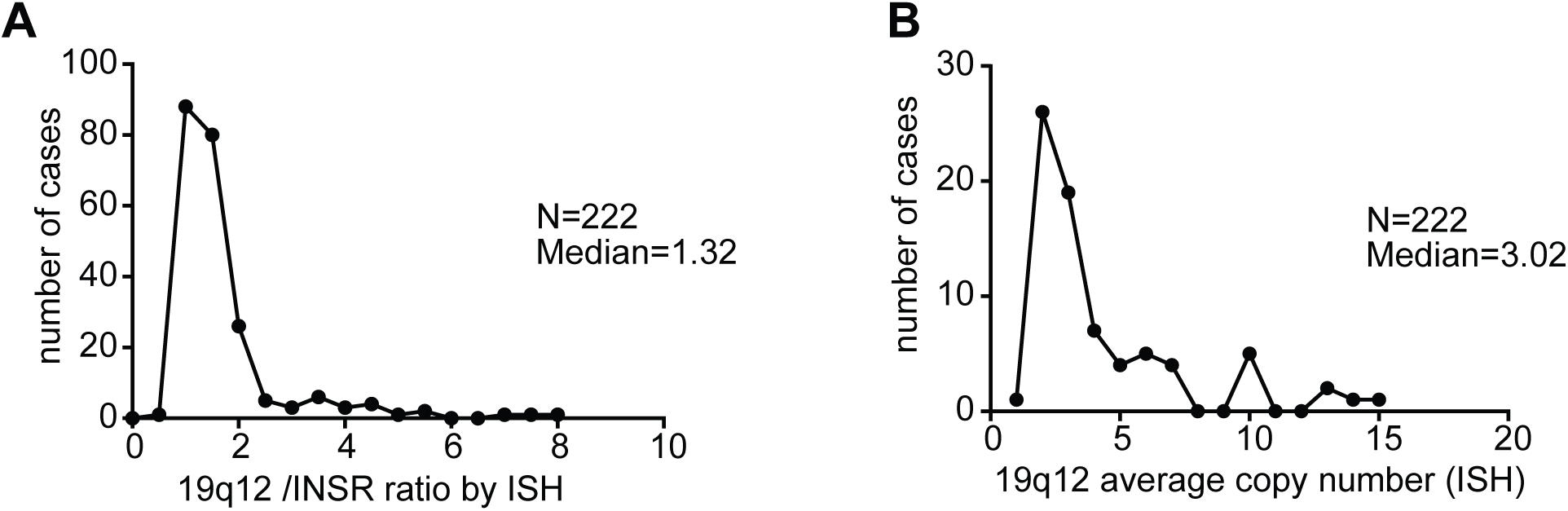
Distribution of 19q12 ISH scores in breast cancer cases

**Supplementary Figure S2:**
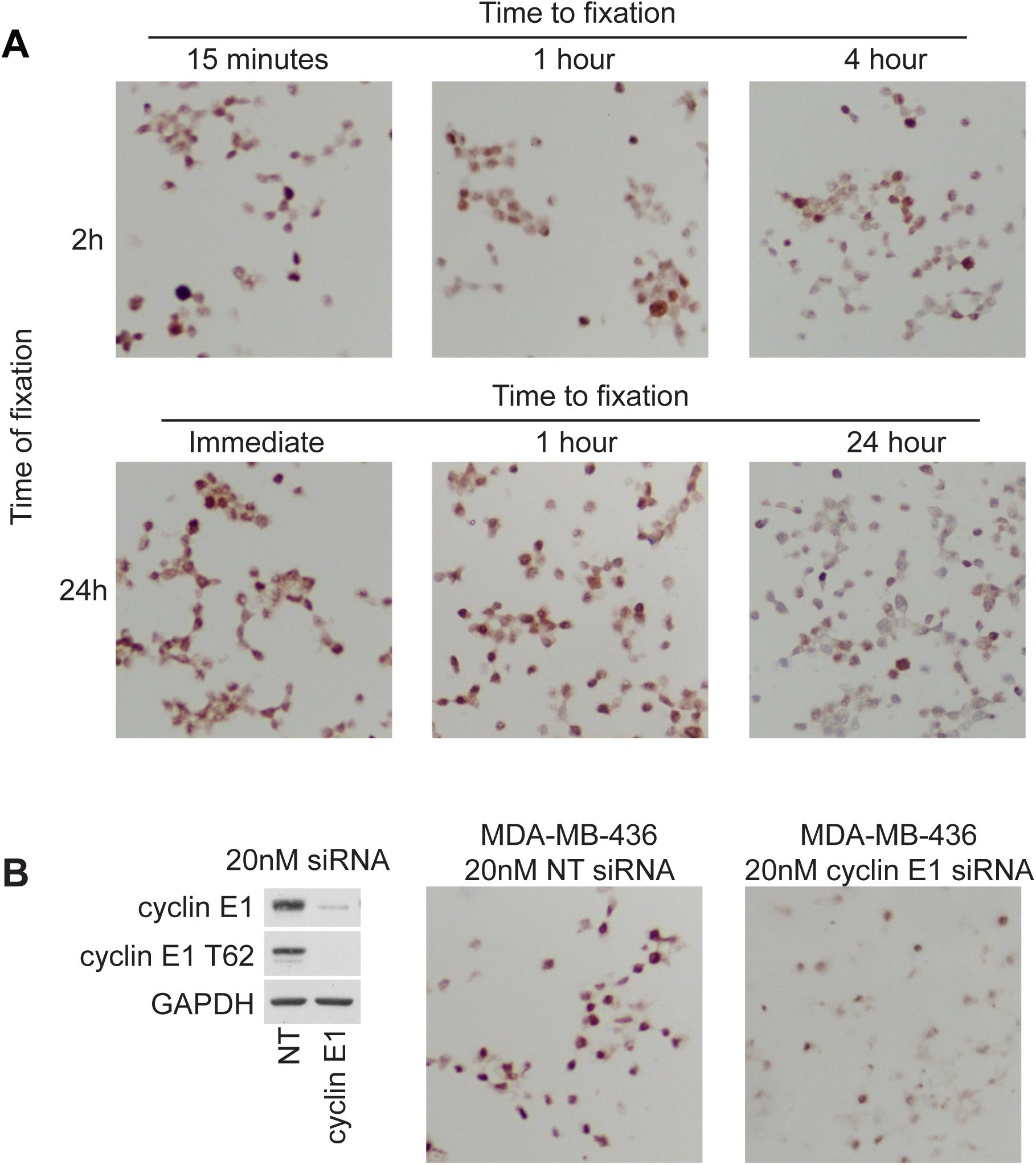
Phospho cyclin E1 T62 antibody optimisation

**Supplementary Figure S3:**
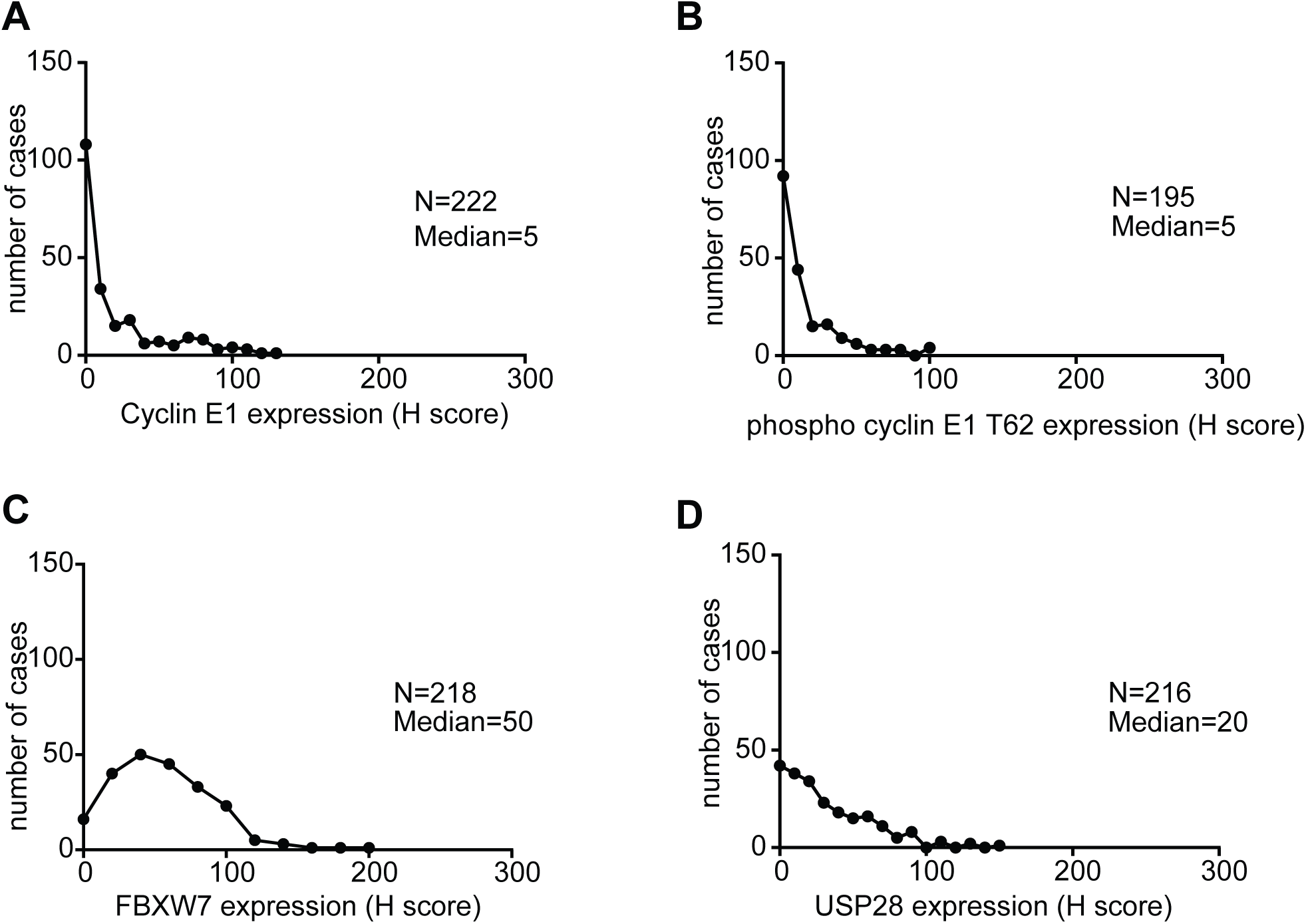
Distribution of H scores in breast cancer cases

**Supplementary Figure S4:**
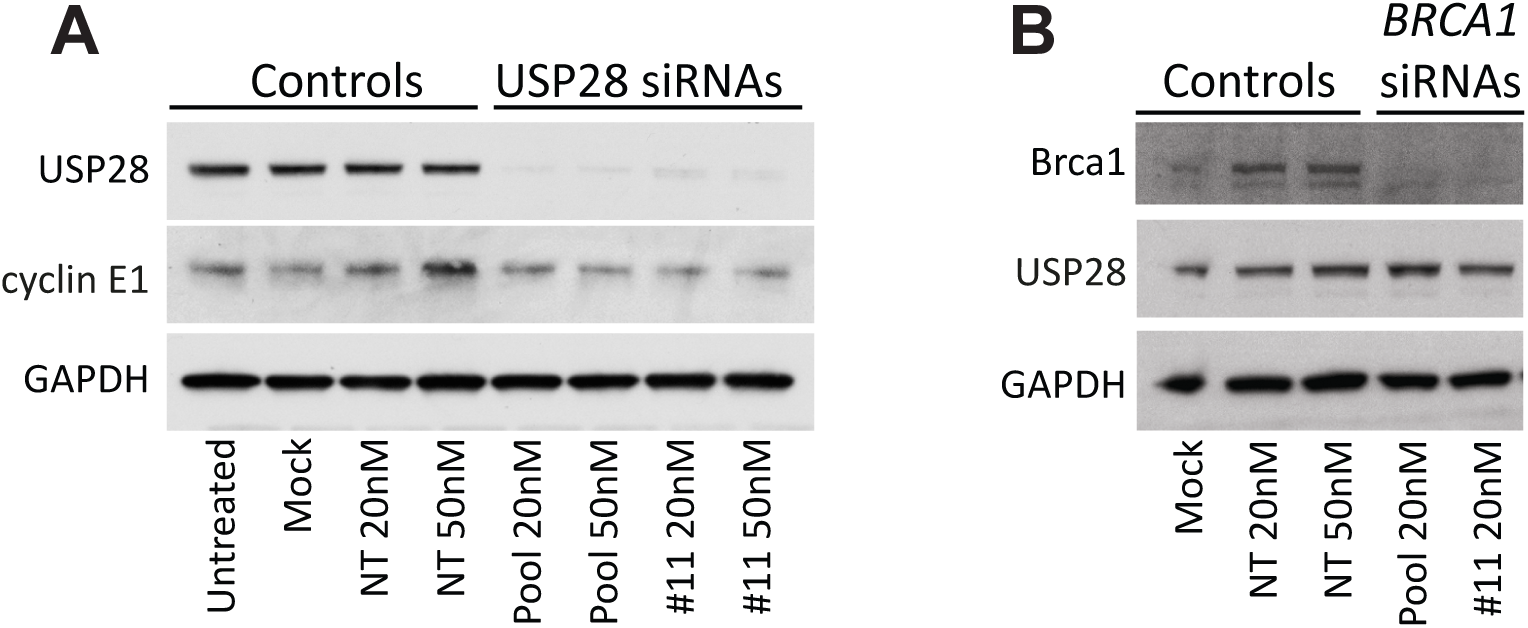
*BRCA1* siRNA does not alter expression of USP28

**Supplementary Figure S5:**
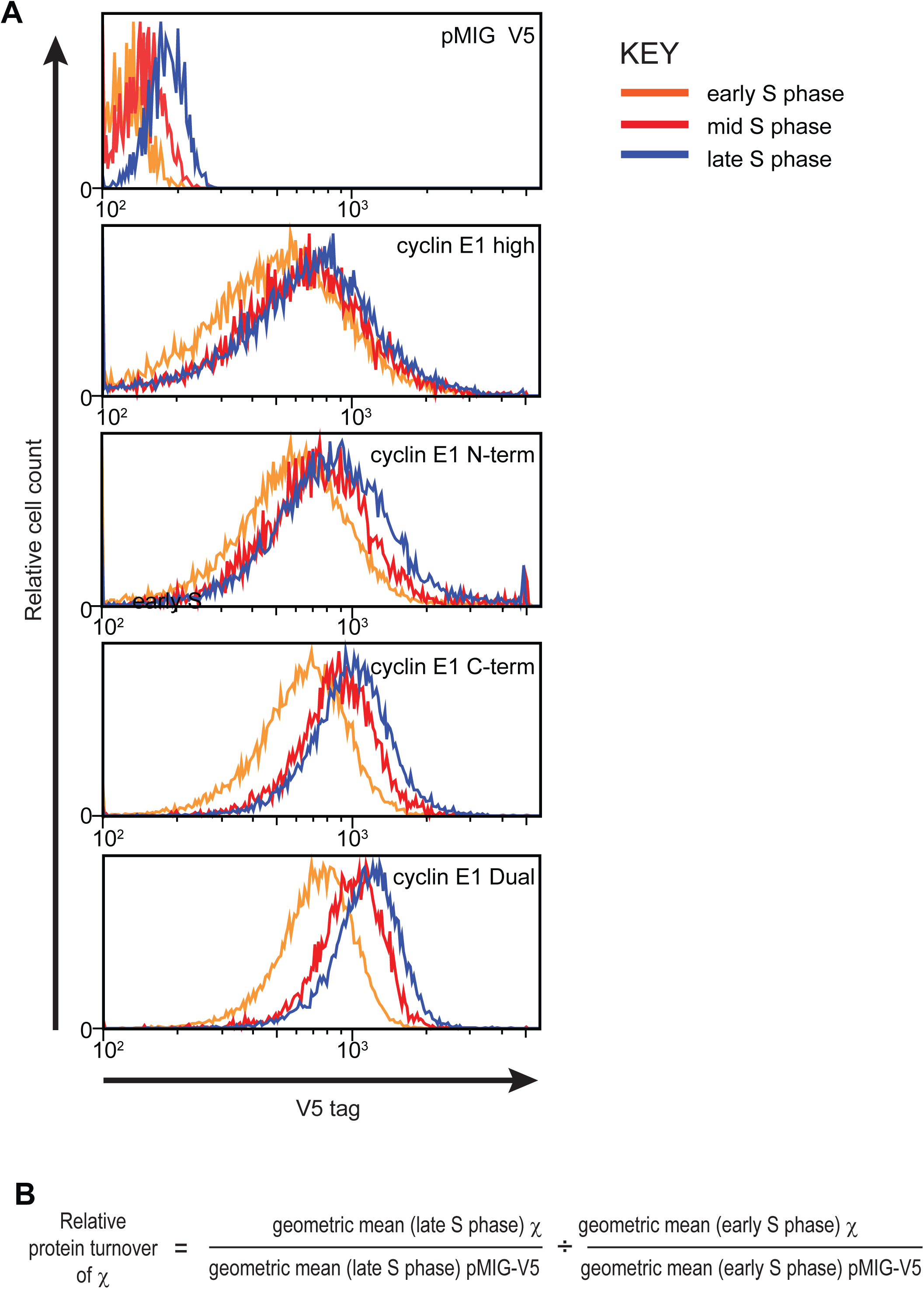
Stability of cyclin E1 constructs in S phase

**Supplementary Figure S6:**
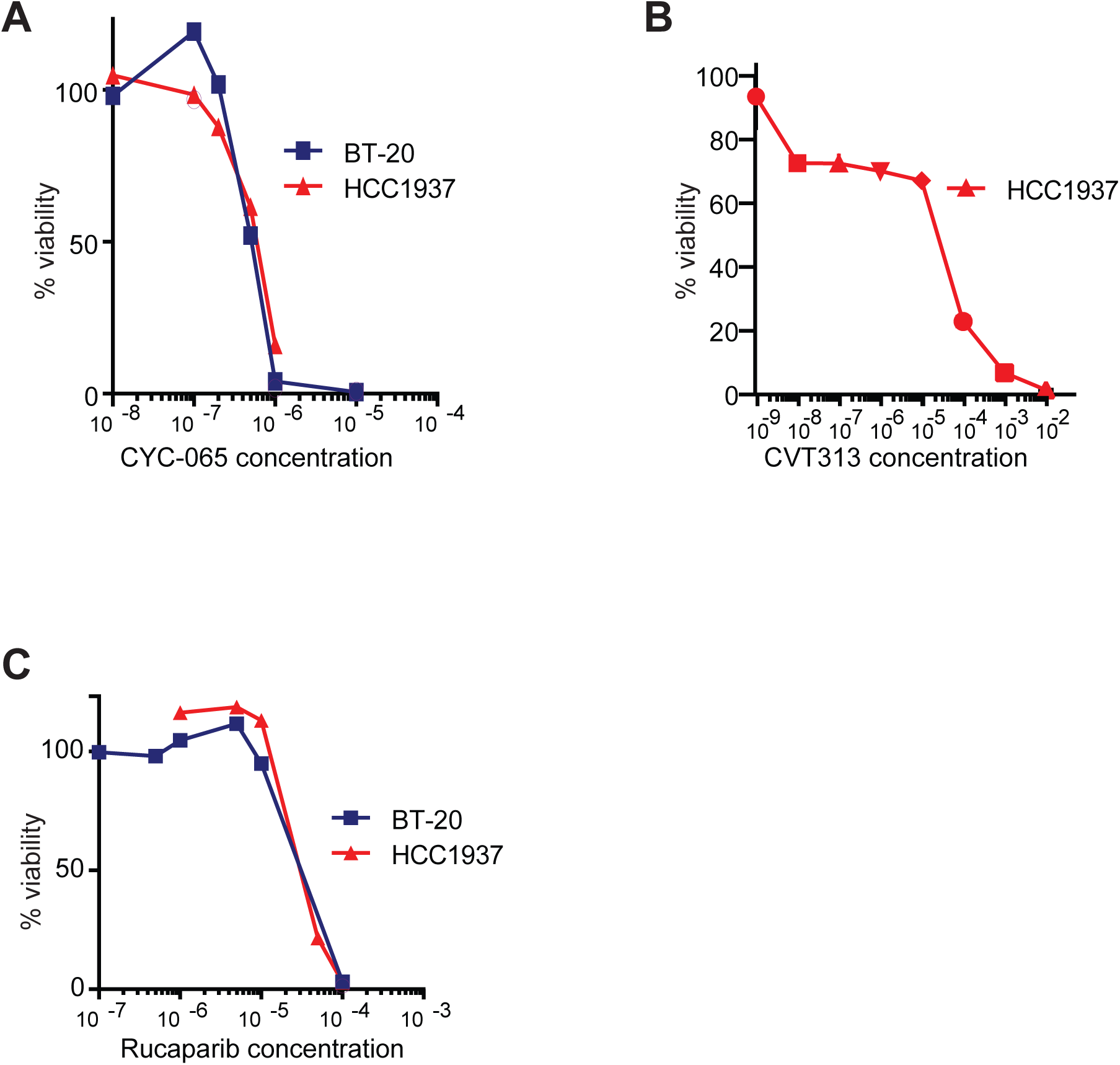
Dose-response curves for cell viability assessment in the BLBC cell lines BT20 and HCC1937

