## Supplementary methods and tables for "Cyclin E1 protein is stabilized in *BRCA1* mutated breast cancers leading to synergy between CDK2 and PARP inhibitors"

**SUPPLEMENTARY INFORMATION**

***in situ* hybridization and cut point determination**

4μm sections from formalin-fixed, paraffin-embedded TMAs were deparaffinized, treated with Cell Conditioning 2 for four cycles of 12 minutes each, followed by ISH protease 3 for 24 min. After co-denaturation, 19q12 DNP and INSR DIG probes were hybridized at 47 °C for 6 hours and washed for 3 cycles at 68 °C for 8 minutes. 19q12 DNP and INSR DIG signals were detected using VENTANA ultraView SISH DNP and VENTANA ultraView RED ISH DIG detection kits, respectively, and counterstained with hematoxylin II and bluing reagent. For image display, the brightness and contrast of each image was adjusted using Photoshop (CS6). 19q12/INSR ratio ≥ 3 and/or 19q12 average number ≥6 were deemed as amplified (1).

Cut-off thresholds for 19q12 amplification were guided by our previously optimised 19q12 ISH assay (1). Amplification status was calculated as 19q12/INSR ratio ≥ 3 and/or 19q12 average number ≥6. The distributions of 19q12/INSR ratio and 19q12 average number are provided in supplementary Figure S1A and S1B respectively.

**T62 antibody validation**

The phospho cyclin E1 (T62) polyclonal antibody (Cell Signaling) was the only IHC grade phospho-cyclin E1 antibody available. As phospho-epitopes are potentially labile, we confirmed that cyclin E1 T62 was detectable in formalin fixed paraffin embedded tissue with a range of fixation conditions. We used paraffin embedded cell blocks of HCT116 colorectal cancer cell lines that were variably fixed in formalin at variable conditions. Different fixation conditions included both variable time to fixation, between pellet stage and 4% paraformaldehyde (PFA) addition; and variable time of fixation, the duration for which the pellet was fixed in 4% PFA. Sections from all variably formalin fixed paraffin embedded cell blocks were stained using the optimized protocol and revealed acceptable robustness of detection (Supplementary Figure S2A).

To further assess the reliability of the p T62 antibody, we tested specificity of the T62 antibody detection by examining cyclin E1 siRNA treated MDA-MB-436 cells. Western blot revealed that cyclin E1 and cyclin E1 T62 expression is lower in cyclin E1 siRNA knocked down cells, which should lead to reduced cyclin E1 T62 expression. Sections from formalin fixed paraffin embedded blocks of each of the cyclin E1 siRNA and control treated cells were stained with the optimized protocol and revealed aligned protein expression to those seen in western blot (Supplementary Figure S2B).

As those experiments revealed the stability, reliability and specificity of T62 antibody, the antibody was next used to stain sections from TMAs of samples of patients enrolled in the KConfab cohort.

**H score and cut point determination for immunohistochemistry**

The H-score was calculated by adding 3 x % of strongly staining (3+) nuclei to 2 x % of moderately staining (2+) nuclei and 1 x % of weakly staining (1+) nuclei, giving a range of 0 to 300. The overall distribution of H score of cyclin E1 expression in all cases (N=222), ranged between 0 and 125 with a median of 5 (Supplementary Figure S3A). The cutoff between high and low cyclin E1 H score was determined by two factors: (1) the previously reported frequency of high cyclin E1 expression in breast cancer and (2) the best association with outcome (minimal p value, Supplementary Table 1), leading to the selection of an H score cut-off of 45. The distribution of mean H scores for phospho-cyclin E1 T62, FBXW7 and USP28 are shown in Supplementary Figure S3B, S3C and S3D respectively. The median H scores were 5 (range: 0 - 100) for cyclin E1 T62 (N=195), 50 (range: 0 - 200) for FBXW7 (N=218) and 20 (range: 0 - 145) for USP28 (N=216). Exhaustion of tissue cores led to unequal numbers of assessable cases for each antibody.

**Flow cytometry for cell cycle specific expression of cyclin E1 and V5**

Cells were incubated overnight at 4°C with antibodies to the E-cyclins (E1: EP435E (Epitomics) or V5 (Invitrogen)) followed by 1 h incubation at room temperature with secondary antibodies (allophycocyanin conjugated goat anti-mouse, fluorescein 5-isothiocyanate conjugated goat anti-rabbit, Jackson Immunoresearch), co-stained with 10μg/mL PI (Sigma) for 2–5h, and incubated with 50μg/mL RNase A (Sigma). Flow cytometry was performed on a FACSCanto (BD Biosciences). Data were analyzed using FlowJo (2). Cells were separated into early, mid and late S phase by identifying the G_1_ and G_2_/M peaks and then partitioning the intervening S phase. Each S phase partition was analysed for expression of cyclin E1 or the V5 tag, where signal intensity was calculated by obtaining the geometric mean signal per cell in gated regions (3,4). Cyclin E1 turnover was calculated as the ratio of expression of late S phase/early S phase. V5-tagged protein turnover was calculated as the ratio of expression of late S phase/early S phase normalized to V5 expression in the pMIG control cell line.

4.	Flow Cytometry: Principles and Applications. Macy MG, editor. Totowa, New Jersey: Humana Press; 2007. 290 p.

**SUPPLEMENTARY FIGURE LEGENDS**

**Supplementary Figure S1**: **Distribution of 19q12 ISH scores in breast cancer cases**

**A:** The distribution of 19q12/INSR ratio by ISH. **B:** The distribution of 19q12 average number.

**Supplementary Figure S2**: **Phospho cyclin E1 T62 antibody optimisation**

**A :** Phospho cyclin E1 T62 expression in HCT116 cell lines at different times to fixations (TTF) and time of fixation (TOF). **B:** Phospho cyclin E1 T62 expression in MDA-MB-436 cells treated with non-targeting control siRNA and cyclin E1 siRNA.

**Supplementary Figure S3**: **Distribution of IHC H scores in breast cancer cases**

**A:** The distribution of mean IHC H scores for cyclin E1 expression in all breast cancer cases. **B:** The distribution of mean H scores for cyclin E1 T62 expression. **C:** The distribution of mean H scores for FBXW7 expression. **D:** The distribution of mean H scores for USP28 expression.

**Supplementary Figure S4: *BRCA1* siRNA does not alter expression of USP28**

**A:** USP28 siRNA was transfected into MDA-MB-468 cells, and lysates collected after 48h and western blotted for USP28, cyclin E1 and GAPDH. **B:** *BRCA1* siRNA was transfected into MDA-MB-468 cells, and lysates collected after 48h and western blotted for Brca1, USP28 and GAPDH.

**Supplementary Figure S5:** **Stability of cyclin E1 constructs in S phase**

**A:** Cells expressing each of the cyclin E1 constructs (pMIG, N-term, C-term, Dual) were analysed by flow cytometry for DNA content (propidium iodide) and V5 (using anti-V5 antibody). Cells were partitioned into early, middle and late S-phase using propidium iodide expression, and V5 expression is shown (x-axis).

**Supplementary Figure S6: Dose-response curves for cell viability assessment in the BLBC cell lines BT20 and HCC1937.**

**A:** Cells were treated with a range of doses of CYC065 for 5 days and relative Alamar Blue staining measured. **B:** Cells were treated with a range of doses of CVT313 for 5 days and relative Alamar Blue staining measured. **B:** Cells were treated with a range of doses of rucaparib for 5 days and relative Alamar Blue staining measured. Experiments performed in triplicate.

**Supplementary Table 1: Determination of cyclin E1 cut-off based on frequency and minimum p-value**

| **H score cut-off** | **% cyclin E1 high** | **OS P value** | **OS HR** | **CI of HR** |
| --- | --- | --- | --- | --- |
| 40 | 42.7 | 0.067 | 0.866 | 0.365-2.054 |
| 45 | 40 | 0.040 | 0.866 | 0.365-2.054 |
| 50 | 36 | 0.063 | 0.548 | 0.233-1.289 |
| 55 | 33.3 | 0.055 | 0.548 | 0.233-1.289 |
| 60/65 | 29.3 | 0.198 | 0.866 | 0.365-2.054 |
| 70 | 26.7 | 0.101 | 0.866 | 0.365-2.054 |

| **All cases (222)** | **19q12 amplified (30)** | | **19q12 non amplified (192)** | |
| --- | --- | --- | --- | --- |
|  | **Cyclin E1≥45** | **Cyclin E1<45** | **Cyclin E1≥45** | **Cyclin E1<45** |
| ***BRCA1* mutated (101)** | 9 (8.9%) | 13 (12.9%) | 24 (23.8%) | 55 (54.4%) |
| ***BRCA1* non-mutated (121)** | 2 (1.8%) | 6 (5.4%) | 6 (5.4%) | 107 (96.4%) |
| **Total** | 11 | 19 | 30 | 162 |

**Supplementary Table 2: Distribution of *BRCA1* mutated and *BRCA1* non-mutated breast cancer cases versus 19q12 amplification and cyclin E1 expression status**
